# Axonal Mechanotransduction Drives Cytoskeletal Responses to Physiological Mechanical Forces

**DOI:** 10.1101/2025.02.11.637689

**Authors:** Grace L Swaim, Oliver V. Glomb, Yi Xie, Chloe Emerson, Zhuoyuan Li, Daniel Beaudet, Adam G. Hendricks, Shaul Yogev

**Author notes:** These authors contributed equally (Grace L Swaim, Oliver V. Glomb).

## Abstract

Axons experience strong mechanical forces due to animal movement. While these forces serve as sensory cues in mechanosensory neurons, their impact on other neuron types remains poorly defined. Here, we uncover signaling that controls an axonal cytoskeletal response to external physiological forces and plays a key role in axonal integrity. Live imaging of microtubules at single-polymer resolution in a *C. elegans* motor neuron reveals local oscillatory movements that fine-tune polymer positioning. Combining cell-specific chemogenetic silencing with targeted degradation alleles to distinguish neuron-intrinsic from extrinsic regulators of these movements, we find that they are driven by muscle contractions and require the mechanosensitive protein Talin, the small GTPase RhoA, and actomyosin activity in the axon. Genetic perturbation of the axon’s ability to buffer tension by disrupting the spectrin-based membrane-associated skeleton leads to RhoA hyperactivation, actomyosin relocalization to foci at microtubule ends, and converts local oscillations into processive bidirectional movements. This results in large gaps between microtubules, disrupting coverage of the axon and leading to its breakage and degeneration. Notably, hyperpolarizing muscle or degrading components of the mechanotransduction signaling pathway in the axon rescues cytoskeletal defects in spectrin-deficient axons. These results identify mechanisms of an axonal cytoskeletal response to physiological forces and highlight the importance of force-buffering and mechanotransduction signaling for axonal integrity.

## Introduction

Maintenance of axonal integrity necessitates mechanisms that buffer and respond to a range of mechanical forces. Axons are subject to internal forces generated by the cytoskeleton and molecular motors, or to external forces such as tissue growth, impact shocks or, particularly at the periphery, animal movement and muscle contraction^1–3^. While recent evidence points to force-buffering mechanisms in the axon, whether and how the axonal cytoskeleton can actively respond and adjust to physiological mechanical forces is unknown.

The Membrane Periodic Skeleton (MPS), a periodic lattice of spectrin tetramers and actin filaments, is a major force-buffering element in the axon^4,5^. Spectrin is held under tension along the longitudinal axis of the axon, where it acts as a shock absorber, likely thanks to its ability to extend under stretch^6–9^. The MPS is also important for radial contractility of the axon and for anchoring channels and signaling molecules^10–15^. Consistent with these multiple functions, pathogenic spectrin mutations lead to a wide range of neurological disorders, making it challenging to precisely pinpoint causal events that lead from spectrin dysfunction to degeneration^16–19^. Cytoskeletal abnormalities have been observed in spectrin mutant neurons from flies and mice, and in *C. elegans* axons break and degenerate in *β-spectrin/unc-70* mutants during locomotion^20–22^. However, how these cellular defects arise following loss of spectrin has not been identified.

In non-neuronal cells, the small GTPase RhoA is a major hub for mechanotransduction. RhoA integrates mechanical signals from integrins and cadherins to regulate actomyosin contractility, signaling and gene expression^23,24^. In neurons, RhoA has mostly been studied in the context of its negative effect on axon growth and regeneration^25,26^. Conversely, the roles of RhoA and actomyosin contractility during homeostasis in uninjured mature axons have been less thoroughly investigated.

Here we found that mechanotransduction signaling, controlled by RhoA and spectrin, regulates the distribution of axonal microtubule (MT) polymers in response to forces exerted on the axon by muscle contractility. Axonal MTs are organized as a partially overlapping linear array, thus providing continuous coverage of the axon^27–29^. While many mechanisms for stabilizing and immobilizing this array have been described, whether MTs can adjust their positioning in adult axons to ensure array continuity is unknown. Using live imaging, neuron-specific temporally controlled degradation alleles and chemogenetic silencing of muscle, we found that muscle contractility leads to local back-and-forth oscillations of axonal microtubules that depend on RhoA, Talin and non-muscle myosin II. Reducing the tension-buffering capacity of the axon by degrading β-spectrin or preventing its axonal delivery caused ectopic RhoA activation and actomyosin redistribution in the axon and drove large-scale MT movements, leading to the formation of gaps between MTs. These phenotypes could be suppressed by inhibiting muscle contractility, indicating that they represent an exaggerated mechanotransduction response. Furthermore, RhoA degradation suppressed breakage of axons lacking β-spectrin, underscoring the contribution of this pathway to cellular dysfunction in spectrin mutants.

## Results

### UNC-76/FEZ1, UNC-69/SCOC and UNC-70/β-spectrin are required for axonal microtubule distribution

To study the axonal cytoskeleton in its native environment, we visualized microtubules (MTs) in DA9, a bipolar cholinergic motor neuron in *C. elegans*. The DA9 axon emerges from the cell body in the preanal ganglion, extends a commissure to the dorsal cord, and grows anteriorly (**Figure 1A**)^30^. MT organization in DA9 is stereotypic, with 4-7 polymers at each cross-section along the proximal axon, and 1-3 MTs in cross sections of the distal axon (defined here as the distal 50% of the axon after the turn of the commissure)^31^. Similar MT densities – and stereotypic transitions in density – are found in axons from invertebrates to mammals^32–34^. MTs in DA9 are reliably visualized with two validated transgenes: GFP::TBA-1/α-tubulin (which labels the entire polymer) and RFP::PTRN-1/CAMSAP, which labels the minus-end (**Figure 1A**)^31,35^.

**Figure 1:**
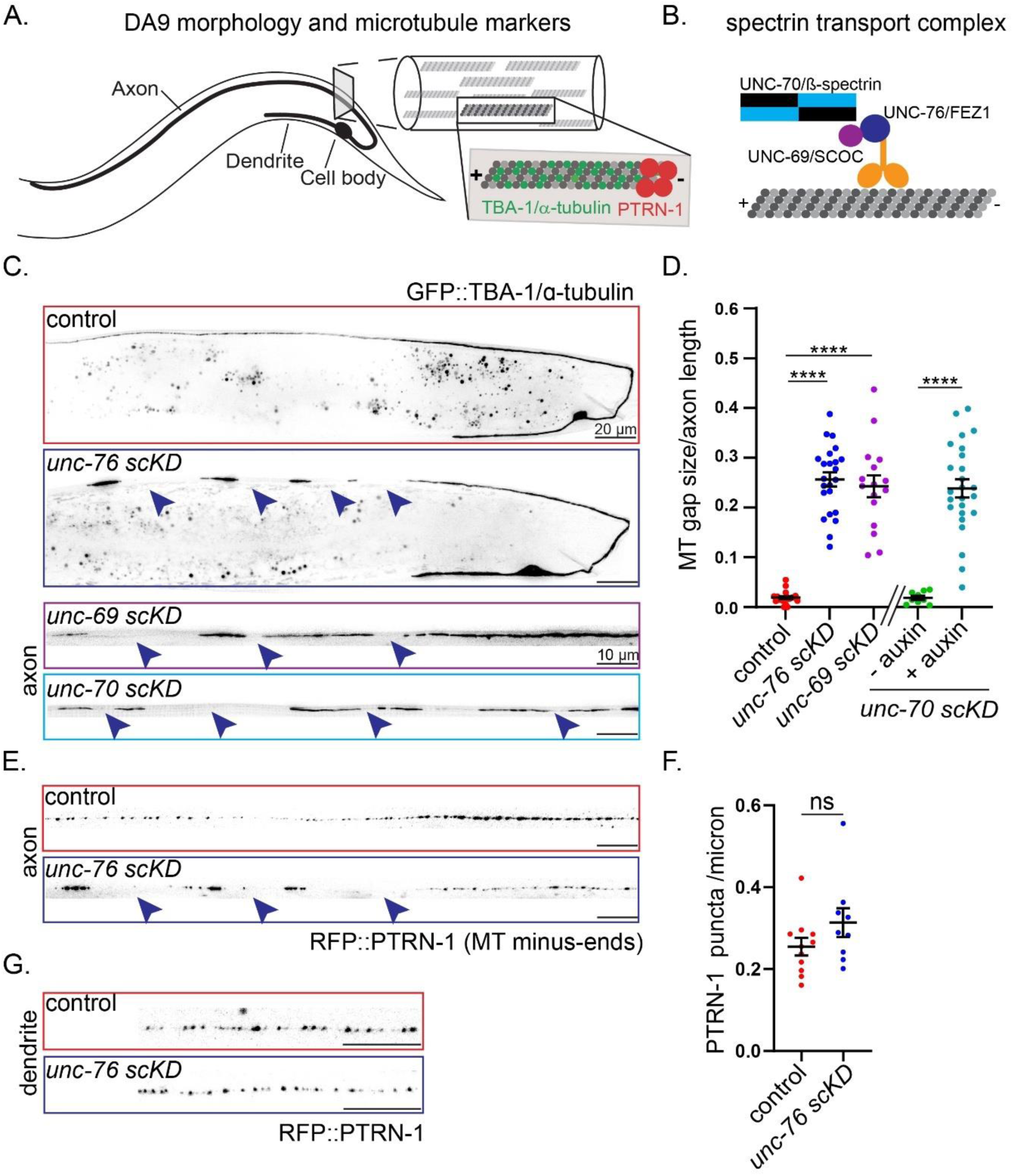
UNC-76/FEZ1, UNC-69/SCOC and UNC-70/b-spectrin are required for axonal microtubule distribution. (A) Diagram of the cholinergic motor neuron DA9, showing microtubule (MT organization) and the markers used for MT lattice (GFP::TBA-1/a-tubulin) and MT minus-ends (RFP::PTRN-1). (B) schematic representation of the spectrin transport complex, consisting of UNC-76/FEZ1 and UNC-69/SCOC. (C) MT organization (GFP::TBA-1) in control, *unc-76 scKD*, *unc-69 scKD*, or *unc-70 scKD* mutants. Control and *unc-76 scKD* image shows the overall structure of DA9, whereas *unc-69 scKD* and *unc-70 scKD* are line scans of the axon. In these images and all subsequent figures, animals are oriented with their anterior to the left, and thus the DA9 axon is oriented from proximal right to distal left. Note gaps in MT continuity (arrowheads) in mutants. Scale bar is 20 microns for the images of the whole DA9 neuron and 10 microns for the line scans. (D). Quantification of C. Note that *unc-70 scKD* utilizes the TIR1/AID degradation system, and so control is animals not raised on auxin, which is required for the degradation. Kruskal-Wallis Test with Dunn’s multiple comparisons (*unc-76 scKD* or *unc-69 scKD* vs control). Unpaired T test (*unc-70 scKD* with or without auxin treatment). **** indicates p<0.0001. n=9-25, error bars show standard error of the mean (SEM). (E) Representative images showing the distribution of minus-ends (RFP::PTRN-1) in control and *unc-76 scKD* mutants. (F) Quantification of RFP::PTRN-1 puncta in control and *unc-76 scKD* mutants showing that despite their misdistribution, the overall number of MT polymers is not affected by degradation of UNC-76. Mann-Whitney test, n=9-11, error bars show SEM. (G) Representative images of dendrites from control and *unc-76 scKD* mutants showing no changes in MT distribution.

In a candidate screen for regulators of axonal MT organization, we found that a mutation in the kinesin-1 adaptor and putative MT binding protein UNC-76/FEZ1 (*unc-76(e911))* disrupts the axonal pattern of GFP::TBA-1/α-tubulin, yielding regions that are devoid of polymers, which we call MT gaps (**Figure S1**). *unc-76* mutants show axon navigation, growth and fasciculation defects^36,37^, which introduce variability into the analysis of MT phenotypes. We therefore induced degradation of UNC-76 specifically in DA9 using a CRISPR-engineered allele (*unc-76 scKD*, see Methods for strain details and degradation protocols) in which axon growth is only mildly impaired and fasciculation and navigation are unaffected^38^. *unc-76 scKD* animals show a drastically altered MT organization pattern, with large gaps and accumulations of polymers between the gaps (**Figure 1C, 1D**). Examination of the minus-end marker RFP::PTRN-1 revealed a similar pattern of gaps (**Figure 1E**). To determine whether MT gaps represent a loss of MT polymers or a defect in their distribution, we estimated MT numbers by counting the number of PTRN-1 puncta (see Methods). We found no significant difference between control and *unc-76 scKD,* suggesting that MT distribution defects underlie the phenotype (**Figure 1F**). These observations indicate that UNC-76 functions cell-autonomously in DA9 to maintain axonal microtubule organization.

The UNC-76 homolog FEZ1 can bind MTs through a C-terminal domain^39^, raising the possibility that UNC-76 functions by directly regulating MT dynamics (i.e. polymer growth and shrinkage). We found that expression of FEZ1 in DA9 robustly suppressed the MT distribution defect of *unc-76 scKD* animals, but the MT binding domain was not required for this rescue (**Figure S1**). This observation suggests that UNC-76 function is conserved, but it does not involve direct MT binding. To further investigate the relationship between MT dynamics and MT distribution defects in *unc-76 scKD* mutants, we asked whether suppressing MT dynamics would influence MT distribution. We examined MT dynamics and gaps in a β-tubulin mutant allele (*tbb-2(qt1)*) which was previously shown to strongly suppress microtubule dynamics in DA9^40,41^. *tbb-2(qt1)* mutants neither displayed MT gaps nor suppressed them in *UNC-76 scKD* animals (**Figure S2**). These results suggest that alterations to MT growth and shrinkage are not causing polymer misdistribution in *unc-76 scKD* mutants.

We recently found that UNC-76/FEZ1, together with the short coiled-coil protein SCOC/UNC-69, acts as a kinesin-1 transport adaptor for axonal delivery of spectrin (**Figure 1B**)^38^. In either *unc-76* or *unc-69* mutants, the spectrin membrane periodic skeleton (MPS) is missing from the distal ∼70% of the DA9 axon^38^. To ask whether a defect in MPS formation or maintenance underlies the MT misdistribution phenotype, we induced degradation of *unc-69/SCOC* and *unc-70/β-spectrin* in DA9. Interestingly, degradation of either protein led to a MT misdistribution defect similar to *unc-76 scKD* (**Figure 1C, 1D**). As with *unc-76,* the function of *unc-69* is conserved, since expression of its human homolog SCOC could rescue MT defects in *unc-69 scKD* worms (**Figure S1**). Consistent with the role of UNC-76 and UNC-69 in transporting spectrin exclusively to the axon, MT misdistribution defects in *unc-69, unc-76* and *unc-70 scKD* mutants were observed exclusively in the axon, while dendritic MTs labeled with RFP::PTRN-1 appeared correctly distributed (**Figure 1G**). Taken together, these results suggest that axonal spectrin is required for normal MT distribution in the axon.

### Reduced oscillations and excessive movements underlie MT misdistribution in axons lacking β-spectrin

To ask how degradation of UNC-70, UNC-76 or UNC-69 leads to MT misdistribution, we carefully examined MT dynamics and movements using live imaging (**Movie S1**). Due to high MT density in axons, end binding proteins (EBs) are usually used to visualize polymer dynamics^42^. However, since EBs only report on the status of growing MT ends, they are not suitable for visualizing whole polymer movements. To visualize the entire MT, we used a GFP::TBA-1/α-tubulin, which we have extensively validated as a reliable MT marker with single-polymer resolution in DA9^31,35^. MT movement can be distinguished from MT growth when both polymer ends change location synchronously. In wildtype animals, along with plus-end growth and shrinkage, we observed that MTs often displayed low amplitude back-and-forth movements (termed here “oscillations”), which to the best of our knowledge have not been previously described for axonal MTs (**Figure 2A**). Oscillations had an amplitude of less than one micron and did not lead to a strong displacement in polymer mass in 1-3 minutes of imaging. Conversely, in *unc-70*, *unc-76* and *unc-69 scKD* conditions, we observed significant polymer movements (**Figure 2C, 2D**). MT movement in the mutants was bidirectional, slow (averaging 0.1-0.2 microns/sec) and relatively processive. Interestingly, MT oscillations were reduced in all three mutants (**Figure 2B**), raising the possibility that the processive MT movements occur at the expense of MT oscillations.

**Figure 2:**
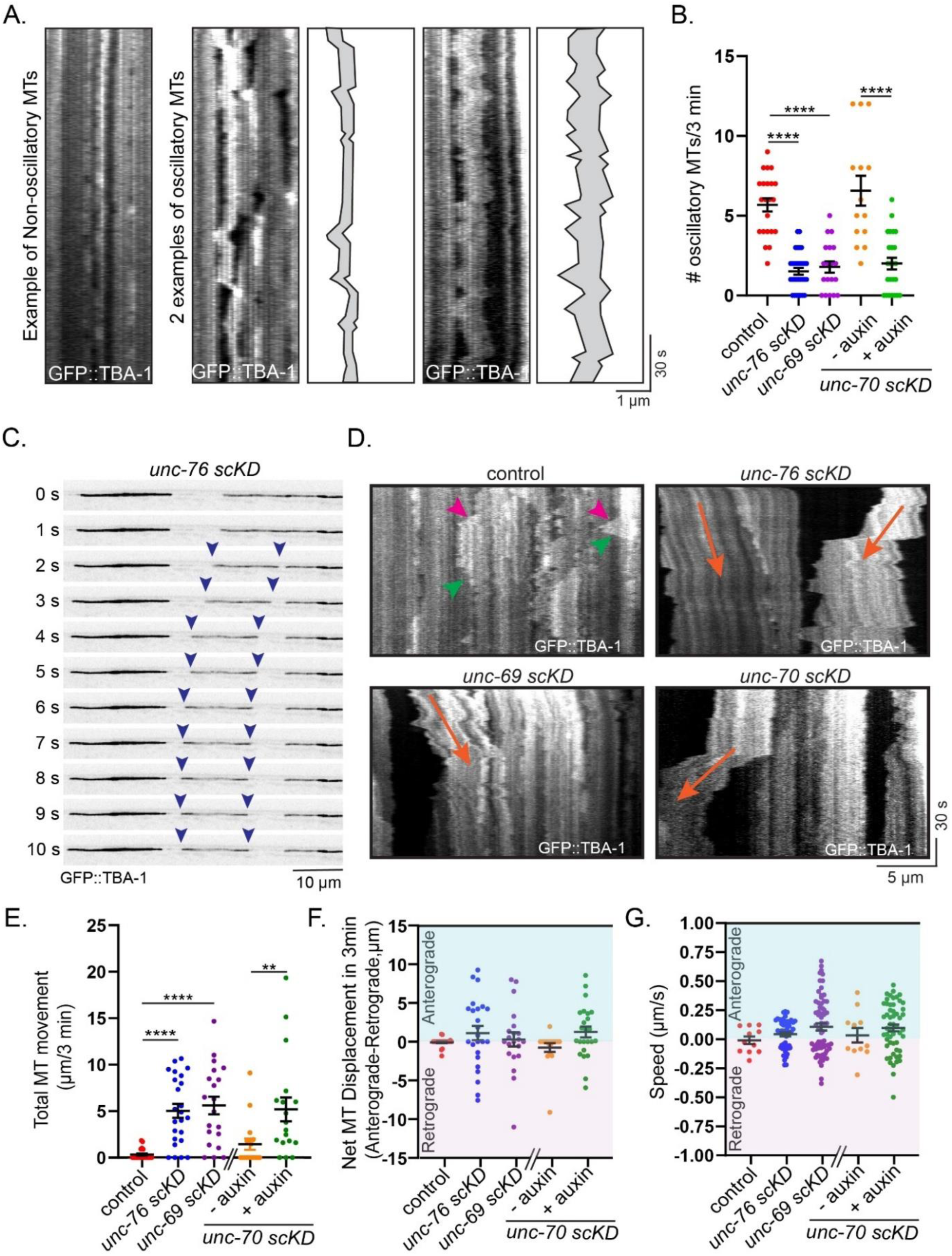
Reduced oscillations and excessive movements underlie MT misdistribution in axons lacking β-spectrin. (A) Representative kymographs showing stationary MTs compared to oscillatory back-and-forth dynamics. Oscillating MTs are highlighted in schematics. The movement of an entire polymer can be distinguished from tip dynamics since both ends of the MT move coordinately. (B) Quantifications of oscillatory microtubules in each genotype. Kruskal-Wallis test with Dunn’s multiple comparison (*unc-76 scKD* or *unc-69 scKD* vs control) or Mann-Whitney Test (*unc-70 scKD* with or without auxin). **** indicates p<0.0001, n=14-31, error bars show SEM. (C) Frames from a timelapse movie showing movement of a single MT in *unc-76 scKD* mutants. Note that the polymer is not moving on other polymers. (D) Representative kymographs highlight MT movements in *unc-76 scKD, unc-69 scKD,* and *unc-70 scKD* mutants compared to control axons. Orange arrowheads highlight long, processive MT movements, magenta and green arrowheads indicate MT plus-end growth and shrinkage, respectively. (E-G) Quantifications of MT displacements in indicated background. n=17-24, error bars show SEM. (E) Shows combined (anterograde and retrograde) movements. Kruskall-Wallis test using Dunn’s multiple comparisons (*unc-76 scKD* or *unc-69 scKD* vs control, **** indicates p<0.0001) and Mann-Whitney (*unc-70 scKD* with or without auxin, ** indicates p=0.0059), error bars show SEM. (F) shows the net displacement in each movie (i.e, the sum of retrograde events is subtracted from the sum of anterograde events). (G) shows average speed per kymograph.

We next tested whether canonical MT motors dynein and kinesin-1 are responsible for the MT movements. We used a temperature sensitive lethal dynein mutant (*dhc-1(or195ts)*)^43^ and an auxin-inducible kinesin-1 degron (*unc-116 scKD*)^38^ to circumvent the requirement for these motors in early development and shifted the animals to the restrictive condition after axon outgrowth (see Methods). Interestingly, both mutants did not significantly affect MT movements in *unc-76 scKD* animals, suggesting that these motors are not required for MT movements (**Figure S3**). We also tested whether MT dynamicity is required for movement by generating *tbb-2(qt1); unc-76 scKD* animals. Reducing MT growth and shrinkage did not affect MT polymer movement, consistent with the observation that it also did not suppress MT gaps at steady state (**Figure S2**). This also confirms that the movements we observe are not due to treadmilling. Taken together, these results suggest that MT misdistribution in mutants for spectrin or its axonal transport adaptors arises because of excessive MT polymer movements.

### Reorganization of actin and non-muscle myosin II in axons lacking spectrin

While imaging MT movements, we often observed polymers moving across gaps (**Figure 2C**), suggesting that MT movement does not require a MT track, consistent with the observation that MT-motors kinesin-1 and dynein were not required for MT movements (**Figure S3**). To identify other cytoskeletal elements that might be involved in MT movements, we visualized F-actin using LifeAct::RFP (**Figure 3A**). In agreement with previous reports^44^, most F-actin visualized with these probes was concentrated in DA9 presynapses in control animals. Conversely, degradation of UNC-70 or UNC-76 led to a striking misdistribution of F-actin to the distal axon (**Figure 3A, B**). Distal F-actin localized preferentially in the gaps between distal MT accumulations and at the edges of accumulations (**Figure 3A**), raising the possibility that it may be involved in their movements. These findings indicate that axonal spectrin is required for F-actin distribution.

**Figure 3:**
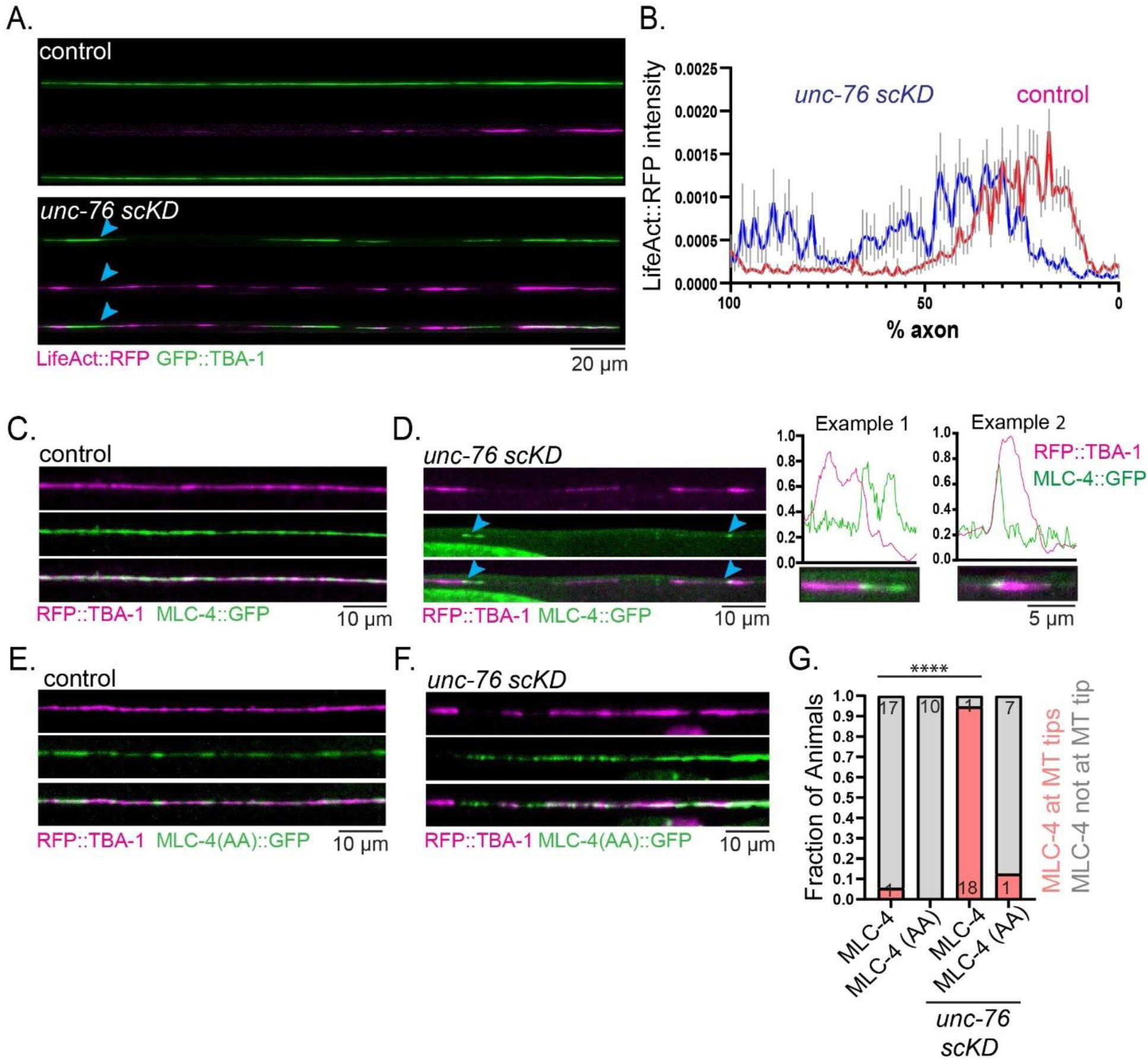
Reorganization of actin and non-muscle myosin II in axons lacking spectrin. (A) Images of tubulin (GFP::TBA-1) and F-actin (LifeAct::RFP) in control and *unc-76 scKD* mutants showing reorganization of actin, with preferential accumulation at the edges of MT patches (arrowhead) Scale bar is 10 μm. (B) Average F-actin signal intensity across the axon in control (n=11) and *unc-76 scKD* mutants (n=10). Distal is left and proximal is right. (C, D) Representative image showing the axonal distribution on non-muscle myosin II regulatory light chain MLC-4 and MTs (GFP::TBA-1). Scale bar is 10 μm. (C) shows control and (D) shows *unc-76 scKD.* Zoomed-in examples show the association of MLC-4 with MT ends in *unc-76 scKD* mutants. Scale bar is 5 μm. (E,F) MLC-4 (AA) appears patchy in control axons (E) and does not relocalize to MT ends in *unc-76 scKD* mutants (F). Scale bar is 10 μm. (G) Fraction of animals with MLC-4 signal localized to the edge of MT patches as shown in panel D in indicated genotypes. MLC-4::GFP localization is compared between control (n=18) and *unc-76 scKD* (n=19) animals, statistical test is Fisher’s exact test, p<0.0001.

To determine whether the misdistribution of F-actin is accompanied by myosin misdistribution, we expressed in DA9 a previously described fusion between GFP and the myosin regulatory light chain MLC-4^45^. In control animals, GFP::MLC-4 was diffusely localized throughout the axon (**Figure 3C**). Degradation of UNC-76 (**Figure 3D**) led to relocalization of GFP::MLC-4, which concentrated around MT tips at the distal axon (**Figure 3D**). GFP::MLC-4(AA), a construct in which two Rho Kinase/ROCK phosphorylation sites that promote myosin II activation were mutated to Alanine^46^, was not as diffuse as GFP::MLC-4 in control axons and did not undergo a localization to MT tips upon degradation of UNC-76 (**Figure 3E, F**). These results suggest that axonal spectrin is required for the distribution of axonal F-actin and myosin and raise the possibility that F-actin and myosin might promote MT misdistribution in axons lacking spectrin.

### Non-muscle myosin II activity is required for microtubule movements and oscillations

To test if F-actin and myosin are required for MT movements, we imaged axonal GFP::TBA-1/α-tubulin dynamics in *unc-76 scKD* animals that were acutely treated with actin and myosin targeting drugs for 5 minutes before imaging (see Methods section). Since the effective concentration of the drug that penetrates the worm’s cuticle is uncertain, some drugs were also tested on *bus-5(br19)* mutants that increase cuticle permeability^47,48^ (see Strain List). In all cases, we confirmed that drug treatments did not cause lethality by documenting MT growth and shrinkage in treated axons. We found that the F-actin stabilizing drug Jasplakinolide (30 μM) effectively blocked F-actin dynamics, but MT movements were unaffected, suggesting that force from actin polymerization is not involved in MT movements (not shown). Conversely, inhibiting non-muscle myosin II with Blebbistatin (32 μm) significantly suppressed MT movements compared to DMSO control (**Figure 4A, B**). ML-7 (0.6 mM), an inhibitor of myosin light-chain kinase also reduced MT movements, but the effect was not statistically significant under these acute conditions (**Figure 4B**). These data suggest that myosin II activity is required for processive MT movements in *unc-76 scKD* axons.

**Figure 4:**
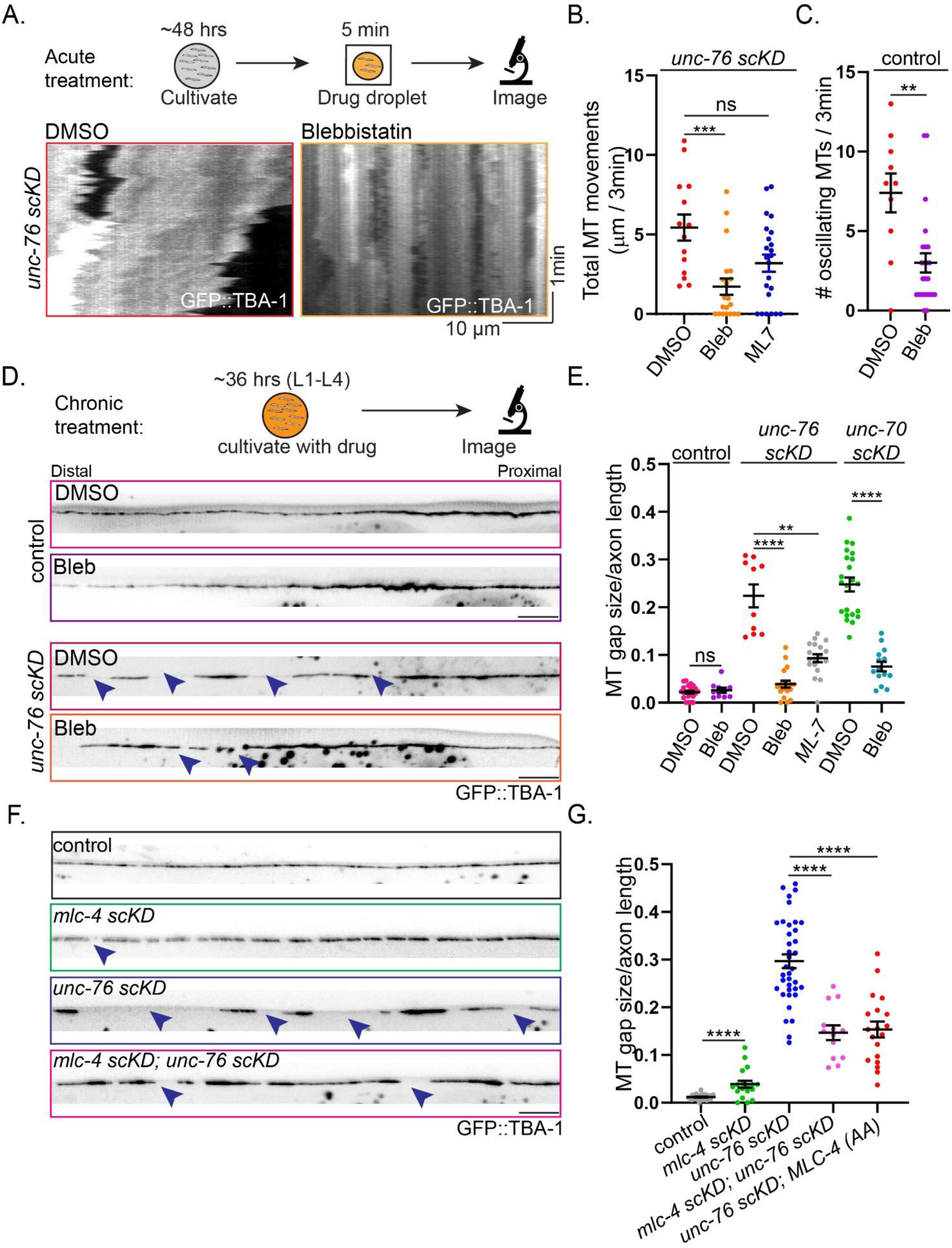
Non-muscle myosin II activity is required for microtubule movements and oscillations. (A) Cartoon schematizes acute pharmacological treatment of animals. Representative kymographs highlight MT movements in *unc-76 scKD* mutants acutely treated with DMSO or Blebbistatin. (B) Quantification of MT movements in Blebbistatin treated *unc-76 scKD* mutants. n=14-23. Kruskall-Wallis test using Dunn’s multiple comparisons ***=adjusted p value of 0.0007, error bars show SEM. (C) Quantification of oscillatory MTs in control animals, treated with DMSO or Blebbistatin. n=10-24. Unpaired Student’s T test, p=0.0011, error bars show SEM. (D) Cartoon schematizes chronic drug treatments. Animals are transferred onto drug-containing plates after the axon reaches its target as L1 larvae. Axons from control and *unc-76 scKD* mutants raised on DMSO or Blebbistatin plates. Arrowheads highlight gaps in the continuous GFP::TBA-1 signal. Scale bar=10 µm. (E) Quantification of D. Statistical test for control is Mann-Whitney test p=0.7413, error bars show SEM. Statistical test for *unc-76 scKD* is Kruskall-Wallis test using Dunn’s multiple comparisons test with DMSO treatment as the control column, ** p=0.0056, **** p<0.0001, error bars show SEM. Statistical test for *unc-70 scKD* is unpaired Student’s t-test, **** p<0.0001. n=10-22, error bars show SEM. (F) Images of distal axons from the indicated genotypes, expressing GFP::TBA-1/α-tubulin. Control is *mlc-4 scKD* without Auxin. Arrowheads indicate MT gaps. Scale bar=10 µm. (G) Quantification of MT gaps normalized to length of the axon. Statistical test comparing control to *mlc-4 scKD* single mutants is unpaired Student’s t-test, n=18-26, p<0.0001, error bars show SEM. Statistical test for *unc-76 scKD*, *mlc-4 scKD*; *unc-76 scKD*, and *unc-76 scKD*; MLC-4(AA)::GFP is ordinary one-way ANOVA with Dunnett’s multiple comparisons with *unc-76 scKD* as the control column, n=12-23, p<0.0001, error bars show SEM.

Since we observed that the appearance of MT movements in *unc-76 scKD* mutants was accompanied by a reduction in MT oscillations, we tested whether MT oscillations were also dependent on non-muscle myosin II. Blebbistatin treatment strongly suppressed MT oscillations in control animals (**Figure 4C**). Hence, both MT movements and MT oscillations require non-muscle myosin II activity.

Next, we asked whether inhibition of MT movements by myosin II targeting drugs could suppress MT gaps at steady state. In agreement with the idea that MT movements underlie the formation of gaps, *unc-76 scKD* animals raised on NGM plates containing Blebbistatin (32 μm) or ML-7 (61 μm) showed reduced MT gap size compared to DMSO control (**Figure 4D, E**). Blebbistatin also suppressed MT gaps in *unc-70 scKD* mutants, confirming that its effect on *unc-76 scKD* mutants was not due to a spectrin-independent function of UNC-76 (**Figure 4E**).

To corroborate these pharmacological results with a genetic approach, we tested how expression of the dominant negative construct MLC-4(AA) affected MT gaps. In addition, we used CRISPR-Cas9 to generate a single-cell degradation allele of MLC-4. MLC-4 was chosen since it is the only non-redundant gene for a myosin II subunit in *C. elegans.* Both approaches led to a strong suppression of MT gaps in *unc-76 scKD* animals **(Figure 4F, G**). Interestingly, degradation of MLC-4 in control axons also led to MT gaps, although these were much smaller than in *unc-76 scKD* (**Figure 4F, G**). We attribute these gaps to myosin’s function in MT oscillations (see Discussion). Taken together, these results suggest that MT gaps and MT movements in axons lacking spectrin, as well as MT oscillations in control animals, are dependent on non-muscle myosin II activity.

### Spectrin restricts axonal RhoA activity

We next sought to identify the mechanism by which spectrin regulates axonal actomyosin and MT distribution. We focused on the small GTPase RhoA (RHO-1 in *C. elegans*), a well-established upstream regulator of actin polymerization and myosin II activation^49–51^. We took the same approach used to uncover the role of non-muscle myosin II: first, we cultured *unc-76 scKD* mutants on plates containing the RhoA inhibitor Rhosin (1 mM). Next, we generated a dominant negative, GDP locked (T17N) RHO-1 construct and expressed it in DA9. Last, we used CRISPR-Cas9 to generate a cell-specific degradation allele of *rho-1* in the background of *unc-70/β-spectrin scKD* mutants, and induced degradation after the axon has reached its target in L1 larvae. In all three cases, we observed a strong reduction in MT gaps (**Figure 5A, B**). Degradation of RHO-1 also reduced MT oscillations in control animals and reduced MT movement in *unc-70 scKD* mutants (**Figure 5C, D**). Although RhoGEF1 is well known to mediate MT-actin crosstalk, *RhoGEF1/rhgf-1* mutants did not phenocopy RHO-1 scKD (not shown). These results suggest that RHO-1/RhoA activity is required for the formation of MT gaps in axons lacking spectrin.

**Figure 5:**
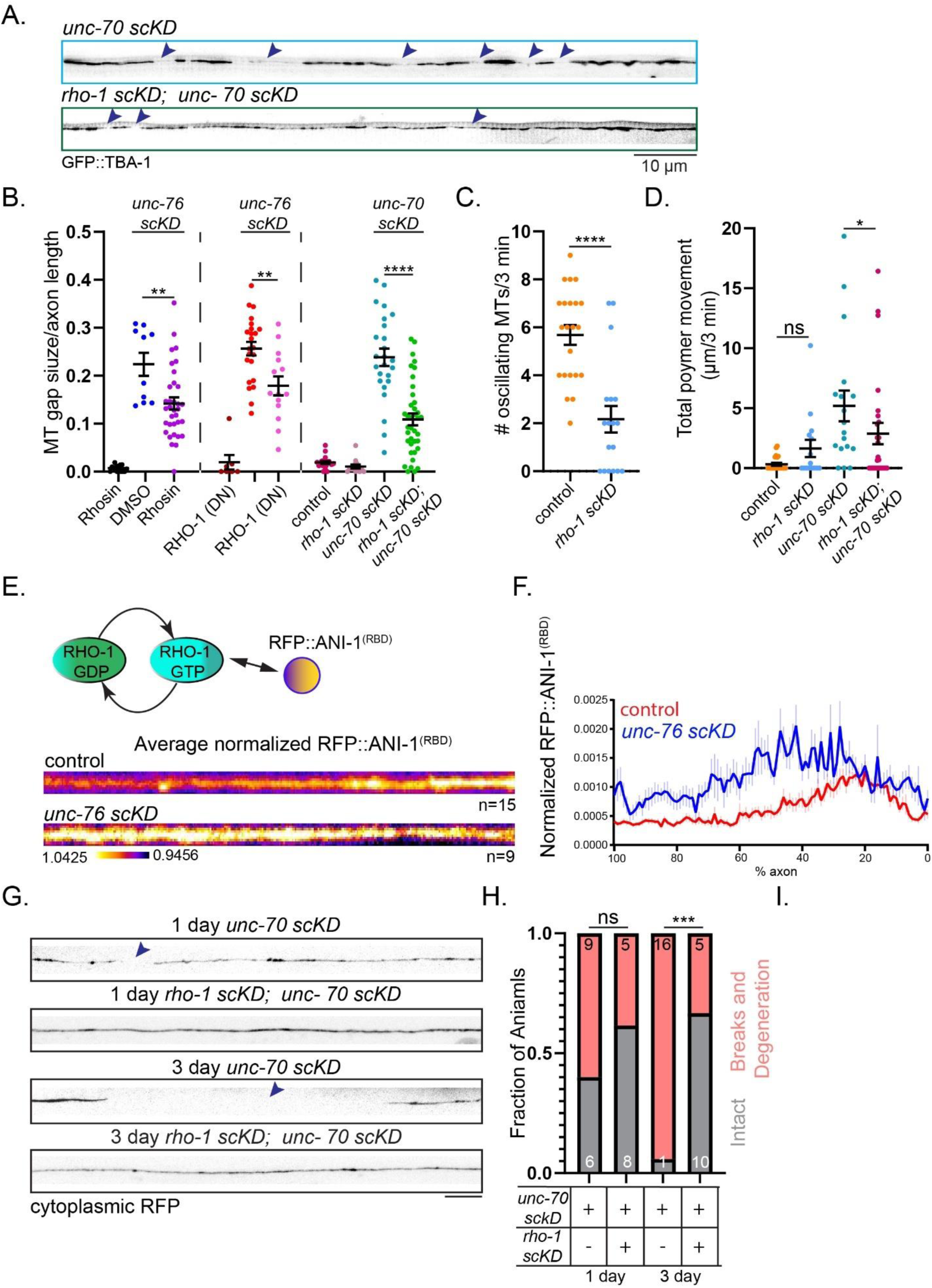
RhoA is inhibited by β-spectrin and is required for axon breakage in β-spectrin mutants. (A) MT distribution in *unc-70 scKD* mutant and *rho-1 scKD*; *unc-70 scKD* double mutants. Degradation of RHO-1 rescues MT gaps (Arrowheads). Scale bar=10 µm. (B) Quantifications of MT gaps in indicated genotypes. Pharmacological inhibition, dominant negative or degradation of RHO-1 reduce MT gaps in axons lacking β-spectrin. DMSO vs Rhosin: Mann Whitney, p=0.0027. *unc-76 scKD* vs *unc-76 scKD;* RHO-1(DN): unpaired Student’s T test, p=0.0026. *unc-70 scKD* vs *rho-1 scKD*; *unc-70 scKD:* unpaired Student’s T test, p<0.0001, n=10-38, error bars show SEM. Control is wildtype (C) Quantification of oscillating MTs in *rho-1 scKD* mutants compared to control. Unpaired t-test, <0.0001, n=15-22, error bars show SEM. (D) Quantification of MT movements in indicated genotypes. Control is wildtype. Statistical test for comparing control to *rho-1 scKD* and *unc-70 scKD* versus *rho-1 scKD; unc-70 scKD* is a Mann-Whitney test, *=p=0.0458 (*unc-70 scKD* vs. *rho-1 scKD, unc-70 scKD*), ns=not significant, n=15-26, error bars show SEM. (E) Schematic shows the design of RhoA activity biosensor. Linescans show averaged pseudo-colored signal intensity of RFP::ANI-1^(RBD)^ in control vs. *unc-76 scKD* mutant animals. In each animal the axonal signal was normalized by the dendritic signal (where UNC-76 degradation does not affect spectrin or MTs) to control for variability in expression. (F) Normalized RFP::ANI-1^(RBD)^ signal intensity in control vs *unc-76 scKD* mutant animals, n=9-15. (G) Axons expressing cytoplasmic RFP show age-dependent breakage (discontinuity indicated by arrowhead) in *unc-70 scKD* mutants that is rescued by co-degradation of RHO-1. Scale bar=10 µm. (H) Quantification of axonal breaks and degeneration. Statistical test is Fisher’s exact test, ns=not significant, p=0.0005. n=13-17.

RHO-1/RhoA activity might be indirectly required for the cytoskeletal rearrangements observed upon spectrin degradation. Alternatively, RHO-1 might undergo excessive activation in mutants lacking axonal spectrin, which would suggest that spectrin is required for controlling RHO-1 activation levels. To distinguish between these possibilities, we expressed an established RhoA biosensor^52^ based on a fragment of Anillin/ANI-1 that binds GTP-RhoA to examine the localization of active RhoA in DA9 (RFP::ANI-1^(RBD)^). We tested several transgenic lines and chose one in which ANI-1 expression did not suppress the MT distribution defect of *unc-76 scKD* mutants (which is expected, given its ability to bind and sequester GTP-bound RhoA). To control for variability in expression, we normalized axonal RFP::ANI-1^(RBD)^ levels to the dendrite, where spectrin remains intact in *unc-76 scKD* mutants^38^. RFP::ANI-1^(RBD)^ was uniformly distributed in control animals, comparable with the distribution of a cytosolic protein. However, in *unc-76 scKD* mutants, the sensor becomes non-uniform and punctate, and its normalized levels increase, particularly in the distal axon (**Figure 5E, F**). These observations suggest that axonal spectrin inhibits RHO-1 activation.

### RHO-1 is required for axon breakage and degeneration of axons lacking spectrin

*unc-70* mutations lead to axon breakage in *C. elegans,* and patient neurons with βII-spectrin mutations show reduced growth and degeneration^17,20^. We previously found that *unc-76 scKD* animals show distal breakage of the DA9 axon^38^. The distal location of these defects is reminiscent of the MT gaps we observe here, raising the possibility that hyperactivation of RHO-1 and the ensuing cytoskeletal rearrangements play a role in the degeneration of axons lacking spectrin. To test this possibility, we generated double mutants *unc-70 scKD; rho-1 scKD* and induced degradation in L1 animals to bypass earlier roles in axon extension. UNC-70 degradation led to axonal breaks and degeneration in ∼60% of 1-day adults and in almost in all 3-day adults. Importantly, co-degradation of RHO-1 led to a rescue of the phenotype in ∼65% of animals at both time points (**Figure 5G, H**), indicating that RHO-1 is required for axon breakage and degeneration of axons lacking spectrin. RHO-1 degradation also allowed *unc-70 scKD* axons in day 3 adults to grow ∼35% longer, from 276.3 ± 23.2 µm to 380.9 ± 24.6 µm (mean ± S.E.M.). These results underscore the major role of RHO-1 in the cellular responses to loss of axonal spectrin.

### Cytoskeletal rearrangements are a mechanotransduction response to muscle contractions and require axonal Talin

Our results thus far suggest that RhoA-dependent MT oscillations are exacerbated in the absence of axonal spectrin. We speculated that MT oscillations in control animals and MT movements in the mutants could represent a mechanotransduction response to tension exerted on the axon because RhoA can be activated in response to mechanical forces, and because spectrin, which is held under tension, acts as a shock absorber for the axon^6,7,23^.

Since muscle contraction exerts tension on the axon, we tested the role of muscle contractility in driving MT gaps in *unc-70 scKD* mutants by crossing them to muscle-specific myosin/*unc-54(e190)* mutants, which are unable to contract their muscles^53^. Whereas *unc-54(e190)* mutants did not display overt MT defects in their axons, the mutation efficiently suppressed the misdistribution of axonal MTs induced by degradation of β-spectrin/UNC-70 (**Figure 6A-C**). These results suggest that muscle contractility is required for cytoskeletal rearrangements in the axon.

**Figure 6:**
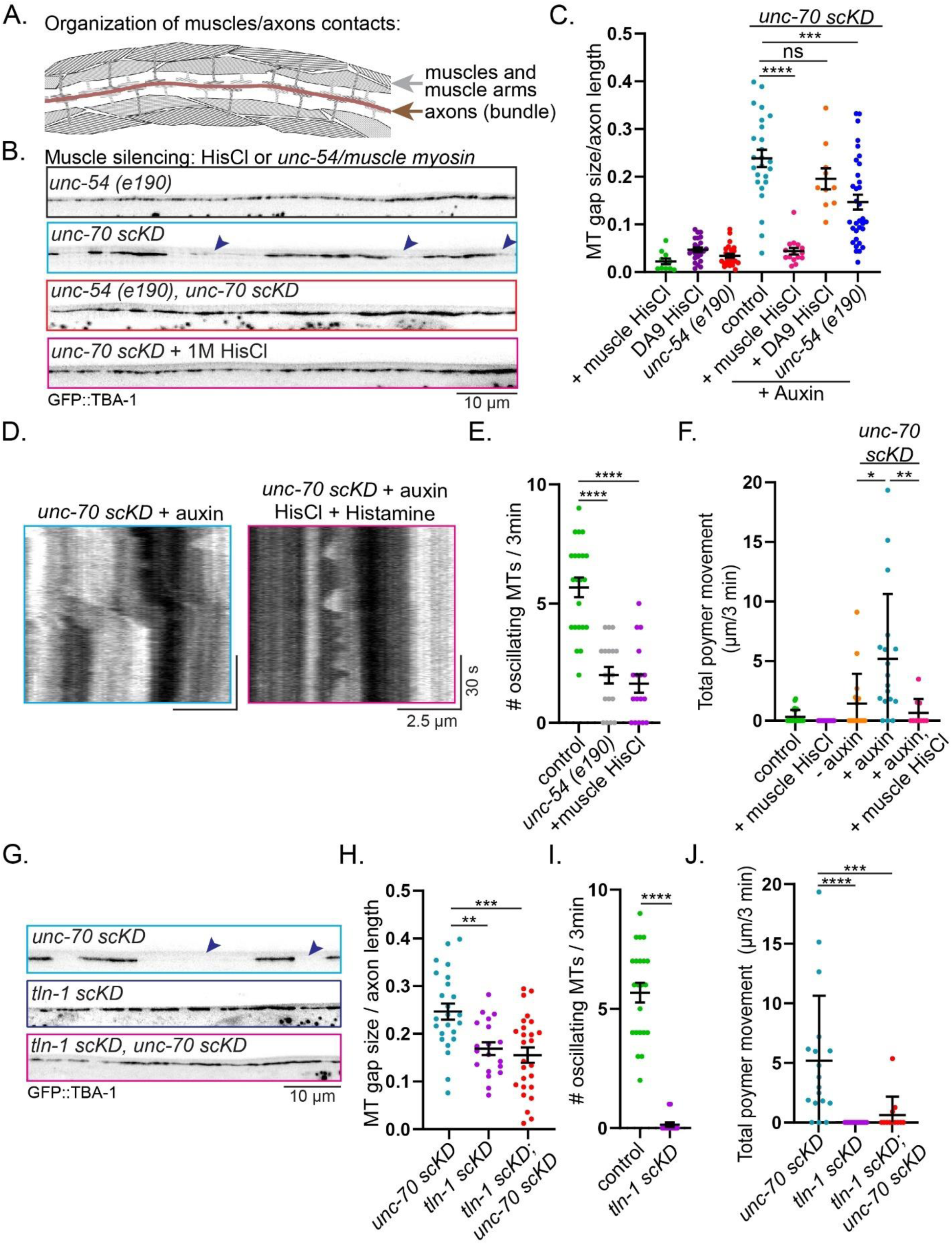
Cytoskeletal rearrangements are a mechanotransduction response to muscle contractions and require axonal Talin. (A) Schematic depicts organization of muscles and muscle arms where they contact axons in the dorsal cord. (B) MT distribution in indicated genotypes. HisCl is a histamine-gated Chloride channel expressed in muscle, which hyper-polarizes the muscle when worms are grown in the presence of Histamine. Worms do not endogenously produce Histamine. Genetically (*unc-54)* or chemogenetically (HisCl) impairing muscle contractility rescues the MT gaps in axons lacking β-spectrin. Scale bar is 10 microns. (C) Quantification of B. Ordinary one-way ANOVA with Dunnett’s multiple comparisons using *unc-70 scKD* as a control column. ****=p<0.0001, ns=not significant, ***=p=0.0001. n=10-33, error bars show SEM. (D) Representative kymographs depicting rescue of MT movement after HisCl treatment in *unc-70 scKD* mutants. (E) Quantifications of the number of oscillating MTs. Hyperpolarizing the muscle strongly reduces the number oscillating MTs in the axon. Kruskall-Wallis test using Dunn’s multiple comparisons test with wildtype as the control column. ****=p<0.0001. n=17-22, error bars show SEM. (F) Quantification of MT movements following muscle silencing. Kruskall-Wallis test using Dunn’s multiple comparisons test with *unc-70 scKD* as the control column. *=p=0.0103, **=p=0.0033. n=10-24, error bars show SEM. (G) Microtubule gaps in *unc-70 scKD* mutants is rescued by co-degradation of TLN-1/Talin in the axon. Scale bar is 10 microns. (H) Quantification of *tln-1 scKD* rescue of *unc-70 scKD* MT gaps. Ordinary one-way ANOVA with Dunnett’s multiple comparisons using *unc-70 scKD* and the control column. **=p=0.0030, ***=p=0.0002, n=19-26, error bars show SEM. (I) Axonal Talin is required for MT oscillations. Mann-Whitney test using wildtype as the control column, ****=p<0.0001, n=14-22, error bars show SEM. (J) Quantification of *tln-1 scKD* rescue of *unc-70 scKD* MT movements. Kruskall-Wallis with Dunnett’s multiple comparisons using *unc-70 scKD* as the control column. **=p=0.0011, ****=p<0.0001, n=12-24, error bars show SEM.

As an orthogonal approach to test the role of muscle activity in regulating the axonal cytoskeleton, we tested whether silencing the muscle through hyperpolarization could similarly suppress cytoskeletal rearrangements in the axon. We expressed a histamine-gated chloride channel (HisCl – note that Histamine is not endogenously produced in *C. elegans*)^54^ specifically in muscles using the *myo-3* promoter, and exposed animals to histamine after axon growth to induce paralysis. Hyperpolarizing the muscle, but not DA9, rescued the distribution of axonal MTs upon degradation of UNC-70/β-spectrin in the neuron. (**Figure 6B, C**). Consistent with the idea that MT gaps are caused by polymer movements, silencing the muscle also reduced MT movements in *unc-70 scKD* mutants (**Figure 6D, F**). These results suggest that cytoskeletal rearrangements induced by degrading spectrin occur in response to muscle contractions.

Next, we asked whether MT oscillations that are observed in control axons are also a response to tension from muscle contraction. This would be consistent with the observation that MT movements in axons lacking spectrin (which require muscle contractility) and MT oscillations in control animals rely on the same mechanisms. *unc-54(e190)* mutants or HisCl expression in muscle led to reduced MT oscillations in the DA9 axon (**Figure 6E**), suggesting that axonal MT oscillations require muscle contraction.

The axonal response to muscle contraction could be mediated by secreted molecules, a mechanically gated channel, or another force-sensing protein. To test the first possibility, we examined whether mutations in *unc-31/CAPS*, which is required for the secretion of dense-core vesicles^55–57^, could suppress MT misdistribution in *unc-76 scKD* mutants. To test the second possibility, we examined mutations in gap junction proteins and mechanically-gated channels that are expressed in DA9^58^. These included *TRPC/trp-1*, *Innexin/unc-7* and *unc-9*, *DegENaC/unc-8*, and *PIEZO/pezo-1*. Neither *unc-31* mutants, nor the mutations in gap junction proteins and channels affected MT distribution in *unc-76 scKD* animals (not shown). While these results do not allow us to definitively rule out a role for vesicle secretion, gap junction proteins and mechanically gated channels, they raise the possibility that other mechanisms might be involved.

We next tested whether the force-sensing and Integrin activating protein Talin/TLN-1 is important for the axonal response to force exerted by muscle contraction. We generated a single-cell degradation allele for TLN-1 with CRISPR-Cas9, and degraded it specifically in DA9 after axon outgrowth. This efficiently suppressed the MT mis-distribution defect in *unc-70 scKD* mutants (**Figure 6G**), suggesting that TLN-1 might be involved in the response of the axonal cytoskeleton to muscle contraction. Similar to degradation of MLC-4, degradation of TLN-1 produced MT gaps in control animals that were smaller than those observed in the absence of axonal spectrin (**Figure 6G, H**). Degradation of TLN-1 also suppressed MT movements in *unc-70 scKD* mutants and suppressed MT oscillations in control animals (**Figure 6I, J**). Together, the results in this section indicate that MT oscillations in control animals, as well as MT movements and gaps in axons lacking spectrin, occur in response to force exerted on the axon by muscle contractions and require axonal Talin.

## Discussion

Axons are continuously exposed to external mechanical forces, yet the axonal response to these forces remains poorly defined. We identified a mechanotransduction pathway that promotes cytoskeletal rearrangements in the axon in response to physiological external forces. In control animals, axonal MTs displayed local oscillations that slightly adjusted their positions. Using cell-specific degradation alleles and chemogenetic silencing of muscles, we found that these oscillations depend on muscle contractility and on the activity of Talin, RhoA and non-muscle myosin II in the axon. Degradation of spectrin or its transport adaptors – which reduces the axon’s ability to buffer tension^7^ – enhanced RhoA activity, driving F-actin and myosin II redistribution and the conversion of local MT oscillations to longer-range processive movements. This led to the formation of large gaps between axonal MTs, axon breakage and degeneration. Importantly, axon breakage and degeneration could be rescued by inhibiting RhoA after initial axon growth. These results reveal a mechanotransduction pathway that regulates cytoskeletal responses in the axon to external forces.

Previous work provides strong evidence that developing axons can respond to externally applied stretch by elongating, increasing MT mass and even adopting axonal fate^59–62^. More recently, abundant transcriptional and proteomic changes were detected in stretched embryonic, but not adolescent, DRG neurons from mice^63^. Our study supports these observations on the ability of the axonal cytoskeleton to remodel in response to external forces. In addition, it uncovers MT oscillations as a previously unrecognized cytoskeletal response to physiological external forces and reveals the mechanotransduction signaling pathway that mediates this response. In previous work, the effects of experimentally stretching developing axons are hypothesized to represent a mechanism that promotes axon elongation *in situ*. In this case, we propose that the role of MT oscillations is to ensure cytoskeletal continuity: dynamically adjusting MT positioning may prevent the formation of gaps during animal movement, when the length of the axon undergoes significant variation in length^6,64^. Consistent with this hypothesis, we observed MT gaps – albeit significantly smaller than in spectrin mutants – in scKD alleles of Talin and MLC-4 (in RhoA scKD the marker is expressed in additional neurons that overlap with the distal DA9, preventing us from reliably detecting gaps). These observations highlight how maintenance of a continuous MT array is finely tuned axons.

The role of spectrin in buffering tension on the axon is well supported by studies *in vivo*, in culture and by modeling^5–7,20,65^. Since the MPS serves additional purposes (i.e. anchoring signaling complexes), spectrin mutations present complex phenotypes that can be challenging to trace back to specific functions. By manipulating muscle contractility in live animals, we provide evidence that cytoskeletal defects in axons lacking *β-spectrin/unc-70* reflect spectrin’s role in buffering tension, suggesting that they are caused by hyperactivated mechanotransduction signaling. This model is consistent with the ectopic activation of RhoA observed in *unc-76 scKD* axons. It is also strongly supported by the suppression MT gaps and movements upon neuron-specific degradation of two proteins that regulate mechanotransduction signaling in non-neuronal cells: RhoA and Talin. We propose that mechanical forces on the axon are transmitted via Talin/ integrin signaling to RhoA. In the absence of tension buffering by spectrin, exaggerated forces experienced by Talin/integrin lead to hyperactivation of RhoA and downstream cytoskeletal remodeling. Interestingly, MT defects in spectrin mutants were noted in the fly NMJ and in Purkinje neurons in mice^21,22^, raising the possibility that the mechanisms we describe here are conserved. It will be informative to determine whether in these cases loss of spectrin also induces cytoskeletal rearrangements through RhoA signaling.

An interesting point that our study did not address is how myosin promotes MT movement. Since we observe MLC-4 accumulating at MT ends, and MLC-4 is required for MT movement and oscillations, we speculate that it is recruited by a protein that binds both MTs and actin. However, the identity of this protein remains to be identified. Non-muscle myosin II is required for MT movements during early neuronal development^66^, also through an unknown linker, raising the possibility that some similarity exists between mechanism that regulate MT distribution in mature and developing axons. An additional point not addressed is the potential contribution of linkers between the MPS and MTs^67–70^ to the observed phenotypes. Our observation of MTs oscillating in control neurons indicate that not all axonal MTs are firmly anchored in place. Furthermore, the ability to suppress MT movements and gaps in spectrin-depleted axon (which lack an MPS by definition) by degrading MLC-4 or RHO-1 suggests that MT immobility can be achieved without cortical anchoring. However, our results cannot rule out that loss of MT anchoring contributes to the MT motility observed in the mutants. Since a range of mechanisms were suggested for MT cortical anchoring, and all the suggested linkers harbor additional unrelated functions, an interesting topic for future studies will be to dissect if and how MT-MPS linkage participates in mechanotransduction regulation of the cytoskeleton.

In summary, this study identifies a cytoskeletal response of non-sensory neurons to physiological forces experienced by their axons and uncovers the mechanotransduction pathway that governs this response.

## Acknowledgements

The authors would like to thank the Yogev lab for discussions and advice. G.S would like to thank B.L, E.S, and K.S for technical assistance. Some strains were provided by the CGC, which is funded by NIH office of Research Infrastructure Programs (P40 OD010440). This project is funded by NIH grant R35GM133573 to S.Y. A.G.H is supported by the Canadian Institutes for Health Research (CIHR PJT-185997). O.V.G was supported by a Walter-Benjamin Scholarship funded by the Deutsche Forschungsgemeinschaft (DFG, German Research Foundation), Project# 465611822.

## Materials and Methods

### Strains and maintenance

Nematode strains were cultured on nematode growth medium plates that were seeded with OP50 bacteria. Animals were grown at 20°C unless indicated otherwise.

### Gene-editing by CRISPR/Cas-9a

All CRISPR based gene-editing experiments were carried out based on a recently updated protocol by the Mello Lab^71^. In brief, in a total injection mix of 20 μl, 0.5 μl Cas9 (*S. pyogenes* Cas9 3NLS, 10μg/μl, IDT# 1081058), 3.7 μg gRNA (1.85 μg/g for each RNA, if two different gRNAs were used), 900fmol dsDNA repair template or 2.2 μg of ssDNA repair template having 32-35bp homology arms and 800 ng PRF4::rol-6(su1006) plasmid was added as a selection marker^72^. Nuclease free water was added to a total volume of 20 μl. The gRNA was synthesized from ssDNA using the EnGen sgRNA Synthesis Kit, *S. pygenes* (NEB# E3322S) and subsequently purifying the gRNA using a Monarch RNA cleanup kit (NEB#T2040). In case of dsDNA as a repair template, the dsDNA was melted and subsequently cooled before adding to the mixture. The mixture was centrifuged at 14.000 rpm for 2 min and 18 μl were transferred to a fresh reaction tube and placed on ice until injected. F1s were genotyped to find heterozygous edited mutants and singled to get homozygotes. Homozygotes were genotyped and Sanger sequenced.

### Plasmids

Plasmids were generated by standard restriction and ligation reactions or by Gibson Assembly and confirmed by Sanger Sequencing. A list of all plasmids that were used or generated in this study is shown in the following table. Sequences and plasmids are available upon request. For each injected construct 2-3 transgenic lines were generated.

### Cell specific expression strategies

We expressed plasmids in DA9 using the promoter of *mig-13. Pmig-13* turns on before animal hatching but labels VA12 (which overlaps with DA9 in the ventral cord) as well as in pharyngeal cells that are required for viability ^73^. To narrow the expression of *Pmig-13* we used an intersectional strategy with a *Punc-4c* promoter driving CRE expression in DA neurons and a LoxP-flanked stop cassette inserted between Pmig-13 and the expressed ORF.

### Temperature-sensitive mutations

For the temperature-sensitive alleles *tbb-2(qt1)* and *dhc-1(or195)*, animals were raised at 15°C and moved to 25°C at the L1 stage and allowed to grow to L4 before being imaged.

### Cell specific degradation

We induced degradation of target proteins using either the ZF1-ZIF-1 or auxin-inducible TIR systems^74,75^. TIR1 and ZIF-1 were expressed using the *mig13* promoter. For auxin-induced degradation, animals were transferred as L1 larvae (after axon outgrowth) from standard NGM plates to plates containing 1 mM Auxin (Alfa Aesar #A10556) and imaged as L4 larvae, unless otherwise noted. Auxin-induced degradation was compared to a control condition that did not induce a knock down. As a control, i) in animals that expressed the TIR1 F-box protein from an extrachromosomal array, animals that lost the extrachromosomal array but co-cultured on auxin plates with their siblings were imaged or ii) in animals that expressed TIR1 from an integrated array, animals were cultured on NGM plates lacking auxin as a control condition.

### Fluorescence microscopy

For live imaging, L4 worms were incubated in 0.5 mM Levamisole diluted in M9 until paralysis and subsequently shifted into a droplet of 0.5 mM Levamisole diluted in M9 on a 10% agarose pad. Still images were captured using a 2% agar pad and incubation in 10mM levamisole in M9 on the pad. Images were acquired with a Laser Safe DMi8 inverted microscope (Leica) equipped with a VT-iSIM system (BioVision) and an ORCA-Flash 4.0 camera (Hamamatsu) controlled by MetaMorph Advanced Confocal Acquisition Software Package. The microscope was equipped with an HC PL APO 63x/1.40NA OIL CS2, a HC PL APO 40x/1.30NA OIL CS2, a HC PL APO 20x/0.8NA AIR and HC PL APO 100x/1.47NA OIL objective as well as 488 nm, 561 nm and 637 nm laser lines.

### Fluorescence quantification

Processing and analysis of raw images and movies was performed using Fiji/ImageJ v2.3.0/1.53f51^76^. Z stacks were assembled by taking multiple images and creating maximum projections. Multiple images were taken if necessary to capture the entire length of DA9 and stitched using the pairwise-stitching plugin with a linear blending fusion method^77^. Time lapse microscopy images with a single channel were stabilized with StackReg using the rigid body transformation method to correct for animal drift during the movie^78^. Alternatively, for imaging conducted in two channels, images were stabilized with HyperStackReg using the affine transformation^79^. To calculate the signal intensity along a neurite, 10pxl thick lines were drawn along the length of the neurite and a signal intensity profile was generated using the plot profile function. To measure microtubule gap sizes, the gaps between GFP::TBA-1 signals were measured using line segments and normalized to the length of the axon.

Representative images shown in figures were adjusted for brightness and contrast. All quantifications were performed on raw images.

### Determining the microtubule and F-actin signal distribution in the axon

To calculate the distribution of GFP::TBA-1 or LifeAct::RFP, fluorescence profiles along the center midline of the axon were acquired. To normalize fluorescence, the signal intensity.at each point on the X axis of the profile was subtracted by the lowest intensity value along the profile, divided by the sum of the signal intensity at each point. Thus, each point had a defined fraction of the total intensity of the profile. Next, intensity values were put in 100 bins, with each bin being 1% of the total length of the line scan. Each bin is the sum of the relative intensity of the pixels it contains. The number of the bin is the position of each point on the X axis and the fraction of the total signal intensity within the bin is its Y axis coordinate.

### Visualizing myosin light chain and microtubule signal distribution in the axon

To highlight the overlapping microtubules and MLC-4 signal in *unc-76 scKD* mutants, fluorescence profiles along the center midline of the axon were acquired. To normalize fluorescence, to normalize fluorescence, the signal intensity at each point on the X axis of the profile was subtracted by the lowest intensity value along the profile, divided by the sum of the signal intensity at each point. Thus, each point had a defined fraction of the total intensity of the profile. The signals are not binned and are presented as per micron.

### Quantifying myosin light chain localization to MT tips

MLC-4::GFP or MLC-4(AA)::GFP signal was defined as localizing to MT tips if specific co-localization to the edges of RFP::TBA-1 patches was observed. Ubiquitous distribution throughout the axon was defined as not at MT tip.

### Estimating microtubule number

Line scans were traced along the axon of animals with RFP::PTRN-1, and the FIJI Find Peaks function was used to identify local maxima^80^. We controlled for axon length by dividing the sum of the peaks by the length of the axon for each line scan.

### Quantifying microtubule dynamics and movement

Kymographs were made using the KymoResliceWide plugin in ImageJ, using the maximum intensity value across width^81^. For presentation purposes, the minimum intensity, brightness, and contrast was modified on kymographs using FIJI. All kymograph analyses were performed before background subtractions.

Microtubule dynamic events were defined as shifts in the GFP::TBA-1 signal of greater than 0.5 microns, where one side of the signal remains stationary and the other shifts.

Microtubule movements were defined as shifts in the GFP::TBA-1 signal where both sides of the signal move, and the movement is consistent in one direction over time.

### Quantifying microtubule oscillations

An oscillating microtubule is defined as a microtubule in a kymograph that undergoes repeated back and forth moment. Oscillating microtubules were counted per kymograph.

### Acute and chronic pharmacological treatment

All drugs were dissolved in DMSO then diluted at least 1:100 in M9 from the initial concentration such that all animals were treated with at most 1% DMSO.

For acute treatments, animals were suspended in a drop of 0.5 mM levamisole+1% DMSO (control) or a drop of 0.5 mM levamisole for 5-30 min until paralyzed, then mounted on a 10% agar pad on a drop of M9 (control) or M9+the drug (concentrations in text). LifeAct was co-imaged with GFP::TBA-1 to visualize MT dynamics and thus ensure that animals were alive during movies. Animals not showing actin dynamics were not quantified, except for animals treated with the actin stabilizing drug Jasplakolinide, which were not quantified if there were no MT dynamics.

For chronic treatments, drugs were pipetted directly onto the surface of 10 mL NGM plates and allowed to permeate the plates for 24 hours. L1 animals were placed on the drug-containing plates, aged to L4, then imaged.

### Drug concentration table

| Drug Name | Acute treatment<br>Concentration | Chronic treatment<br>Concentration | Solvent |
| --- | --- | --- | --- |
| Jasplakinolide | 0.03 mM | NA | DMSO |
| ML-7 | 0.663 mM | 0.0663 mM | DMSO |
| Blebbistatin | 0.03244 mM | 0.03244 mM | DMSO |
| Rhosin | NA | 1mM | DMSO |

### RhoA Biosensor quantification

To calculate the signal intensity of RFP::ANI-1^(RBD)^ along a neurite, 10pxl thick lines were drawn along the length of the neurite and a signal intensity profile was generated using the plot profile function for both the axon and dendrite. The signal intensity of RFP::ANI-1^(RBD)^ at each pixel of the axon was divided by the average dendrite signal intensity to normalize the expression of RFP::ANI-1^(RBD)^ in each genetic background. Next, the signal intensity at each point on the X axis of the profile was subtracted by the lowest intensity value along the profile, divided by the sum of the signal intensity at each point. Thus, each point had a defined fraction of the total intensity of the profile. Then, intensity values were put in 100 bins, with each bin being 1% of the total length of the line scan. Each bin is the sum of the relative intensity of the pixels it contains. The number of the bin is the position of each point on the X axis and the fraction of the total signal intensity within the bin is its Y axis coordinate.

### Quantifying axon breakage and degeneration

Discontinuity in the cytoplasmic RFP signal was interpreted as axon breakage. Animals were counted that had axon breaks.

### Histamine chloride treatment

A plasmid containing the Drosophila histamine-gated chloride channel HisCl1, was expressed under the *mig-13* promoter to silence DA9 or under the *myo-3* promoter to silence muscle cells. L1 animals containing the array were picked onto standard NGM plates. 500uL of 1M Histamine chloride [Histamine dihydrochloride powder, Sigma-Aldrich, CAS 56-92-8] dissolved in M9 was pipetted directly onto the animals in the plates, which were left to air dry overnight. Animals were prepared for imaging as previously described.

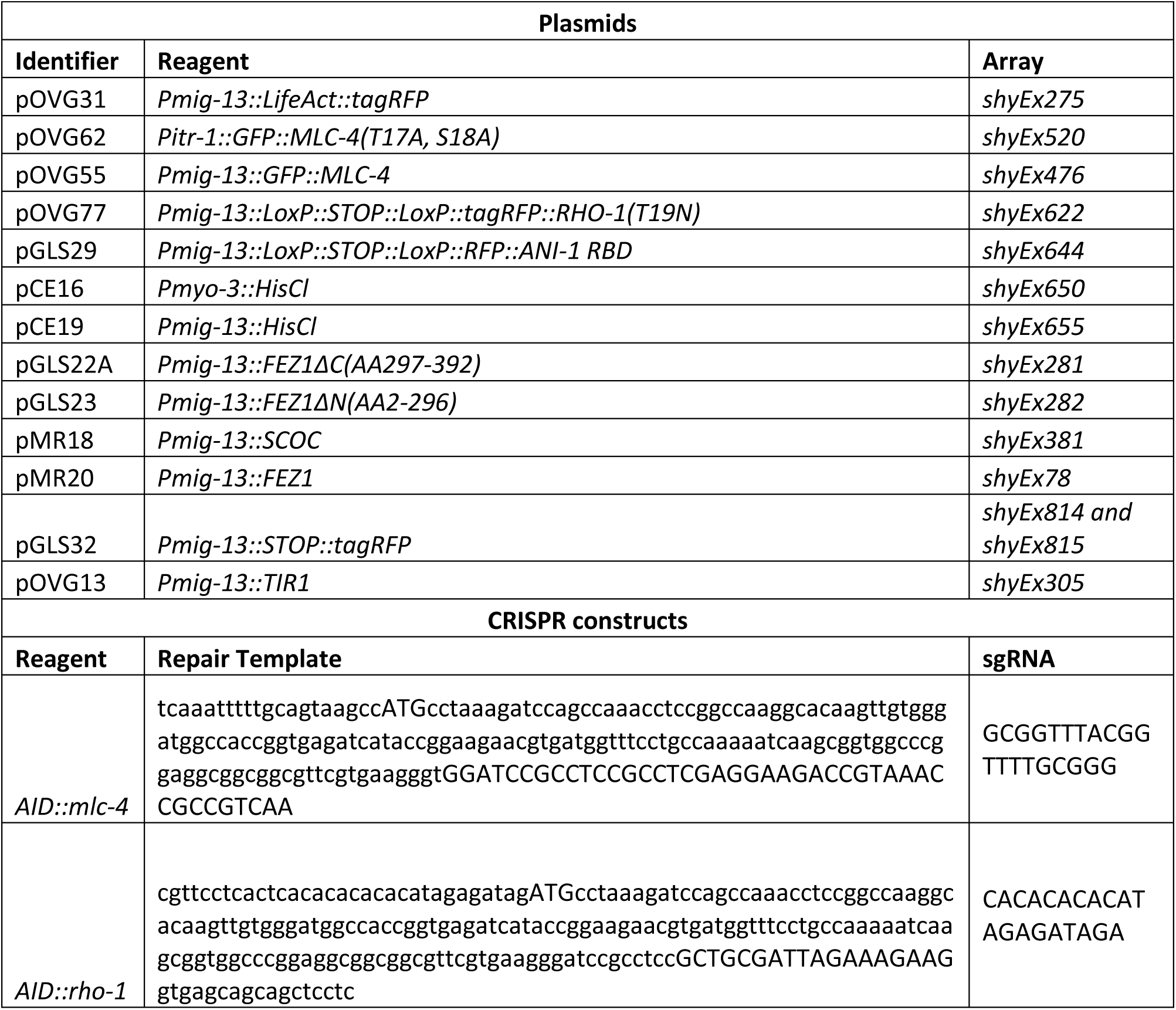
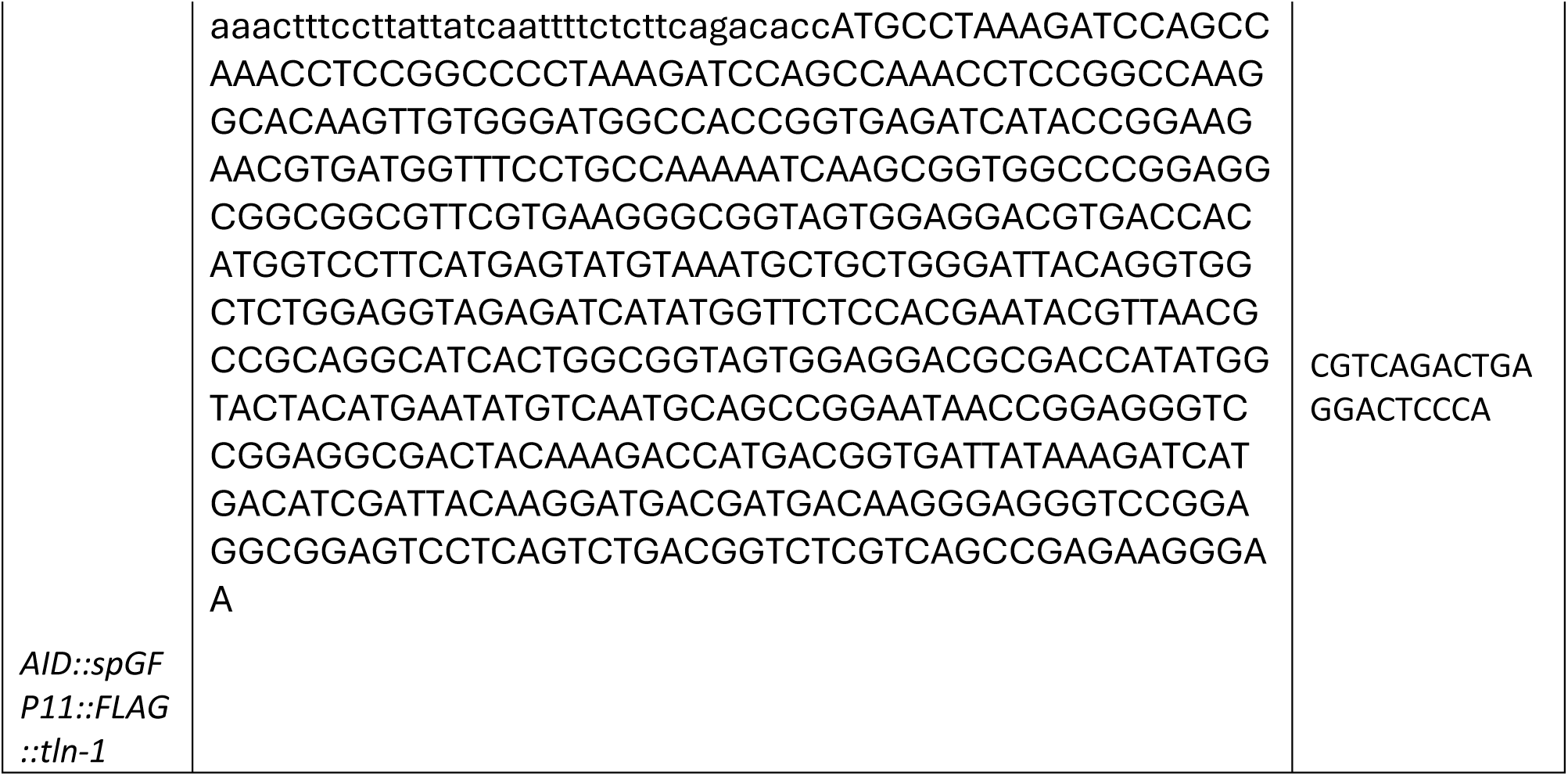

| Strain ID | Genotype |
| --- | --- |
| TV19492 | *tba-1(ok1135) I*; *wySi265[Punc-4C::CRE, Punc-122:RFP] II*; *wyls 463[Pmig-13::LoxP::STOP::LoxP::GFP::TBA-1, Podr-1::RFP]*, *wyls720[Pmig-13::LoxP::STOP::LoxP::TagRFP::PTRN-1, Podr-1::GFP] V* |
| MTS358 | *tba-1(ok1135) I*; *wySi265[Punc-4C::CRE, Punc-122:RFP] II*; *him-8(mn253)*, *shyls9[Pmig-13::ZIF-1, Pttx-3::RFP] IV*; *unc-76(shy19[zf-1::unc-76])*, *wyls463[Pmig-13::LoxP::STOP::LoxP::GFP::TBA-1, Podr-1::RFP] V* |
| MTS837 | *tba-1(ok1135) I*; *wySi265[Punc-4C::CRE, Punc-122:RFP] II*; *unc-69(syb2774[zf-1::7xGFP11::unc-69]) III*; *wyls463[Pmig-13::LoxP::STOP::LoxP::GFP::TBA-1, Podr-1::RFP]*, *shyls16[Pmig-13::ZIF-1, Pttx-3::RFP] V* |
| MTS1369 | *tba-1(ok1135) I*; *wySi265[Punc-4C::CRE, Punc-122:RFP] II*; *unc-70(syb1684[unc-70(1166R + AID + 1167D)]) (V)*, *wyls465[Pmig-13::LoxP::STOP::LoxP::GFP::TBA-1] X*; *shyEx305[Pmig-13::TIR1, Podr-1::RFP]* |
| MTS655 | *tba-1(ok1135) I*, *tbb-2(qt1) I*; *wySi265[Punc-4C::CRE, Punc-122:RFP] II*; *wyls463[Pmig-13::LoxP::STOP::LoxP::GFP::TBA-1, Podr-1::RFP]*, *wyls720[Pmig-13::LoxP::STOP::LoxP::TagRFP::PTRN-1, Podr-1::GFP] V* |
| MTS618 | *tba-1(ok1135) I*, *tbb-2(qt1) I*; *wySi265[Punc-4C::CRE, Punc-122:RFP] II*; *him-8(mn253) IV*; *wyls463[Pmig-13::LoxP::STOP::LoxP::GFP::TBA-1, Podr-1::RFP]*, *shyls16[Pmig-13::ZIF-1, Pttx-3::RFP]*, *unc-76(shy19[zf-1::unc-76]) V* |
| MTS1367 | *tba-1(ok1135) I*; *wySi265[Punc-4C::CRE, Punc-122:RFP] II*; *wyls463[Pmig-13::LoxP::STOP::LoxP::GFP::TBA-1, Podr-1::RFP] V*; *shyEx276[Pmig-13::LifeAct::tagRFP, Podr-1::GFP]* |
| MTS1304 | *tba-1(ok1135) I*; *wySi265[Punc-4C::CRE, Punc-122:RFP] II*; *him-8(mn253) IV*; *wyls463[Pmig-13::LoxP::STOP::LoxP::GFP::TBA-1, Podr-1::RFP]*, *shyls16[Pmig-13::ZIF-1, Pttx-3::RFP]*, *unc-76(shy19[zf-1::unc-76]) V*; *shyEx276[Pmig-13::LifeAct::tagRFP, Podr-1::GFP]* |
| MTS1922 | *tba-1(ok1135) I*; *shyls38[Pmig-13::LoxP::STOP::LoxP::wTagRFP::TBA-1, Podr-1::GFP]*, *wySi265[Punc-4C::CRE, Punc-122:RFP] II*; *shyls9[Pmig-13::ZIF-1, Pttx-3::RFP]*; *shyEx476[Pmig-13::GFP::MLC-4, Podr-1::RFP]* |
| MTS2092 | *tba-1(ok1135) I*; *shyls38[Pmig-13::LoxP::STOP::LoxP::wTagRFP::TBA-1, Podr-1::GFP]*, *wySi265[Punc-4C::CRE, Punc-122:RFP] II*; *shyls16[Pmig-13::ZIF-1, Pttx-3::RFP]*, *unc-76(shy19 [zf-1::unc-76]) V*; *shyEx476[Pmig-13::GFP::MLC-4, Podr-1::RFP]* |
| MTS2102 | *tba-1(ok1135) I*; *shyls38[Pmig-13::LoxP::STOP::LoxP::wTagRFP::TBA-1, Podr-1::GFP]*, *wySi265[Punc-4C::CRE, Punc-122:RFP] II*; *shyls9[Pmig-13::ZIF-1, Pttx-3::RFP]*; *shyEx520[Pitr-1::GFP::MLC-4(T17A, S18A), Podr-1::RFP]* |
| MTS2161 | *tba-1(ok1135) I*; *wySi265[Punc-4C::CRE, Punc-122:RFP] II*; *shyls16[Pmig-13::ZIF-1, Pttx-3::RFP]*, *unc-76(shy19[zf-1::unc-76])*, *wyls463[Pmig-13::LoxP::STOP::LoxP::GFP::TBA-1, Podr-1::RFP] V*; *shyEx520[Pitr-1::GFP::MLC-4(T17A, S18A), Podr-1::RFP]* |
| MTS1481 | *tba-1(ok1135) I*; *wySi265[Punc-4C::CRE, Punc-122:RFP] II*; *him-8(mn253) IV*; *wyls463[Pmig-13::LoxP::STOP::LoxP::GFP::TBA-1, Podr-1::RFP]*, *shyls16[Pmig-13::ZIF-1, Pttx-3::RFP]*, *unc-76(shy19[ZF1::unc-76]) V*; *bus-5(br19) X*; *shyEx276[Pmig-13::LifeAct::tagRFP, Podr-1::GFP]* |
| MTS1507 | *tba-1(ok1135) I*; *wySi265[Punc-4C::CRE, Punc-122:RFP] II*; *wyls463[Pmig-13::LoxP::STOP::LoxP::GFP::TBA-1, Podr-1::RFP] V*; *bus-5(br19) X*; *shyEx276[Pmig-13::LifeAct::tagRFP, Podr-1::GFP]* |
| MTS2209 | *tba-1(ok1135) I; wySi265[Punc-4C::CRE, Punc-122:RFP] II; mlc-4(shy163[AID::mlc-4]); wyls463[Pmig-13::LoxP::STOP::LoxP::GFP::TBA-1, Podr-1::RFP] V; shyEx305[Pmig-13::TIR1, Podr-1::GFP]* |
| MTS2221 | *tba-1(ok1135) I; wySi265[Punc-4C::CRE, Punc-122:RFP] II; mlc-4(shy163[AID::mlc-4]); shyls16[Pmig-13::ZIF-1, Pttx-3::RFP], unc-76(shy19 [zf-1::unc-76], wyls463[Pmig-13::LoxP::STOP::LoxP::GFP::TBA-1, Podr-1::RFP] V; shyEx305[Pmig-13::TIR1, Podr-1::GFP]* |
| MTS3235 | *tba-1(ok1135) I; rho-1(shy193[AID::rho-1]) IV; shyls68 [Pmig-13::GFP::TBA-1, Podr-1::RFP] V; shyEx305[Pmig-13::TIR1, Podr-1::GFP]* |
| MTS2659 | *tba-1(ok1135) I; wySi265[Punc-4C::CRE, Punc-122:RFP] II; rho-1(shy193[AID::rho-1]) IV; shyls52[Pmig-13::TIR1, Podr-1::GFP], unc-70(syb1684[unc-70(1166R + AID + 1167D)]) V; wyls465[Pmig-13::LoxP::STOP::LoxP::GFP::TBA-1, Pmgig-13::RFP] X* |
| MTS2600 | *tba-1(ok1135) I; wySi265[Punc-4C::CRE, Punc-122:RFP] II; him-8(mn253); shyls16[Pmig-13::ZIF-1, Pttx-3::RFP], unc-76(shy19[zf-1::unc-76]), wyls463[Pmig-13::LoxP::STOP::LoxP::GFP::TBA-1, Podr-1::RFP] V; shyEx622[Pmig-13::LoxP::STOP::LoxP::tagRFP::rho-1(T19N), Podr-1::GFP]* |
| MTS3216 | *tba-1(ok1135) I; wySi265[Punc-4C::CRE, Punc-122:RFP] II; him-8(mn253) IV; wyls463[Pmig-13::LoxP::STOP::LoxP::GFP::TBA-1, Podr-1::RFP] V; shyEx622[Pmig-13::LoxP::STOP::LoxP::tagRFP::rho-1(T19N), Podr-1::GFP]* |
| MTS3195 | *tba-1(ok1135) I; wySi265[Punc-4C::CRE, Punc-122:RFP] II; shyls52[Pmig-13::TIR1, Podr-1::GFP], unc-70(syb1684[unc-70(1166R + AID + 1167D)]) V; wyls465[Pmig-13::LoxP::STOP::LoxP::GFP::TBA-1, Podr-1::RFP] X; shyEx814[Pmig-13::LoxP::STOP::LoxP::RFP, Pmyo-2::GFP]* |
| MTS3196 | *tba-1(ok1135) I; wySi265[Punc-4C::CRE, Punc-122:RFP] II; rho-1(shy193); shyls52, unc-70(syb1684[unc-70(1166R + AID + 1167D)]) V; wyls465[Pmig-13::LoxP::STOP::LoxP::GFP::TBA-1] X; shyEx815[Pmig-13::LoxP::STOP::LoxP::RFP, Pmyo-2::GFP]* |
| MTS3215 | *tba-1(ok1135) I; wySi265[Punc4::CRE, Punc-122:RFP] II; wyls463[Pmig-13::LoxP::STOP::LoxP::GFP::TBA-1, Podr-1::RFP] V; shyEx644[Pmig-13::LoxP::STOP::LoxP::RFP::ANI-1 RBD, Pmyo-2::mCherry]* |
| MTS2714 | *tba-1(ok1135) I; wySi265[Punc-4C::CRE, Punc-122:RFP] II; shyls16[Pmig-13::ZIF-1, Pttx-3::RFP], unc-76(shy19[zf-1::unc-76]), wyls463[Pmig-13::LoxP::STOP::LoxP::GFP::TBA-1, Podr-1::RFP] V; shyEx644[Pmig-13::LoxP::STOP::LoxP::RFP::ANI-1 RBD, Pmyo-2::mCherry]* |
| MTS2555 | *tba-1(ok1135) I, unc-54(e190) I; wySi265[Punc-4C::CRE, Punc-122:RFP] II; shyls52[Pmig-13::TIR1, Podr-1::RFP], unc-70(syb1684[unc-70(1166R + AID + 1167D)]) V; wyls465[Pmig-13::LoxP::STOP::LoxP::GFP::TBA-1, Podr-1::RFP] X* |
| MTS2741 | *tba-1(ok1135) I; wySi265[Punc-4C::CRE, Punc-122:RFP] II; shyls52[Pmig-13::TIR1, Podr-1::GFP], unc-70(syb1684[unc-70(1166R + AID + 1167D)]) V; wyls465[Pmig-13::LoxP::STOP::LoxP::GFP::TBA-1, Podr-1::RFP] X; shyEx650[Pmyo-3::HisCl, Pmyo-2::GFP]* |
| MTS2841 | *tba-1(ok1135) I, unc-54(e190) I; wySi265[Punc-4C::CRE, Punc-122:RFP] II; wyls465[Pmig-13::LoxP::STOP::LoxP::GFP::TBA-1, Podr-1::RFP] X* |
| MTS2817 | *tba-1(ok1135) I; wySi265[Punc-4C::CRE, Punc-122:RFP] II; wyls465[Pmig-13::LoxP::STOP::LoxP::GFP::TBA-1, Podr-1::RFP] X; shyEx650[Pmyo-3::HisCl, Pmyo-2::GFP]* |
| MTS2900 | *tba-1(ok1135) I; wySi265[Punc-4C::CRE, Punc-122:RFP] II; shyls52[Pmig-13::TIR1, Podr-1::RFP], unc-70(syb1684[unc-70(1166R + AID + 1167D)]) V; wyls465[Pmig-13::LoxP::STOP::LoxP::GFP::TBA-1, Podr-1::RFP] X; shyEx706[Pmig-13::HisCl, Pmyo-2::GFP]* |
| MTS3203 | *tba-1(ok1135) I, tln-1(shy226[AID::spGFP::FLAG::tln-1]; wySi265[Punc-4C::CRE, Punc-122:RFP] II; shyls52[Pmig-13::TIR1, Podr-1::GFP] V; wyls465[Pmig-13::LoxP::STOP::LoxP::GFP::TBA-1, Podr-1::RFP] X* |
| MTS3035 | *tba-1(ok1135) I, tln-1(shy226[AID::spGFP::FLAG::tln-1]; wySi265[Punc-4C::CRE, Punc-122:RFP] II; shyls52[Pmig-13::TIR1, Podr-1::GFP], unc-70(syb1684[unc-70(1166R + AID + 1167D)]) V; wyls465[Pmig-13::LoxP::STOP::LoxP::GFP::TBA-1, Podr-1::RFP] X* |
| MTS840 | *dhc-1(or195ts), tba-1(ok1135) I; wySi265[Punc-4C::CRE, Punc-122:RFP] II; wyls463[Pmig-13::LoxP::STOP::LoxP::GFP::TBA-1, Podr-1::RFP] V* |
| MTS841 | *dhc-1(or195ts), tba-1(ok1135) I; wySi265[Punc-4C::CRE, Punc-122:RFP] II; wyls463[Pmig-13::LoxP::STOP::LoxP::GFP::TBA-1, Podr-1::RFP], shyls16[Pmig-13::ZIF-1, Pttx-3::RFP], unc-76(shy19[zf-1::unc-76]) V* |
| MTS1610 | *tba-1(ok1135) I; wySi265[Punc-4C::CRE, Punc-122:RFP] II; unc-116(shy92[unc-116::AID]) III; shyls16[Pmig-13::ZIF-1, Pttx-3::RFP], wyls463[Pmig-13::LoxP::STOP::LoxP::GFP::TBA-1, Podr-1::RFP] V; shyEx305[Pmig-13::TIR1, Podr-1::RFP]* |
| MTS2663 | *tba-1(ok1135) I; wySi265[Punc-4C::CRE, Punc-122:RFP] II; unc-116(shy92[unc-116::AID]) III; him-8(mn253) IV; shyls16[Pmig-13::ZIF-1, Pttx-3::RFP], unc-76 (shy19[zf-1::unc-76]), wyls463[Pmig-13::LoxP::STOP::LoxP::GFP::TBA-1, Podr-1::RFP] V; shyEx305[Pmig-13::TIR1, Podr-1::RFP]* |
| MTS364 | *tba-1(ok1135) I; wySi265[Punc-4C::CRE, Punc-122:RFP] II; shyls9[Pmig-13::ZIF-1, Pttx-3::RFP]) IV; unc-76(shy19[zf-1::unc-76]), wyls463[Pmig-13::LoxP::STOP::LoxP::GFP::TBA-1, Podr-1::RFP] V; shyEx78[Pmig-13::FEZ1, Podr-1::GFP]* |
| MTS1120 | *tba-1(ok1135) I; wySi265[Punc-4C::CRE, Punc-122:RFP] II; shyls9[Pmig-13::ZIF-1, Pttx-3::RFP]) IV; unc-76(shy19[zf-1::unc-76]), wyls463[Pmig-13::LoxP::STOP::LoxP::GFP::TBA-1, Podr-1::RFP] V; shyEx281[Pmig-13::FEZ1ΔC(AA297-392), Podr-1::GFP]* |
| MTS1121 | *tba-1(ok1135) I; wySi265[Punc-4C::CRE, Punc-122:RFP] II; shyls9[Pmig-13::ZIF-1, Pttx-3::RFP]) IV; unc-76(shy19[zf-1::unc-76]), wyls463[Pmig-13::LoxP::STOP::LoxP::GFP::TBA-1, Podr-1::RFP] V; shyEx282[Pmig-13::FEZ1ΔN(AA2-296), Podr-1::GFP]* |
| MTS2831 | *tba-1(ok1135) I; wySi265[Punc-4C::CRE, Punc-122:RFP] II; unc-69(syb2774[zf-1::7xGFP11::unc-69]) III; shyls9[Pmig-13::ZIF-1, Pttx-3::RFP]) IV; wyls463[Pmig-13::LoxP::STOP::LoxP::GFP::TBA-1, Podr-1::RFP] V; shyEx381[Pmig-13::SCOC]* |

## Supplementary Material

**Figure S1:**
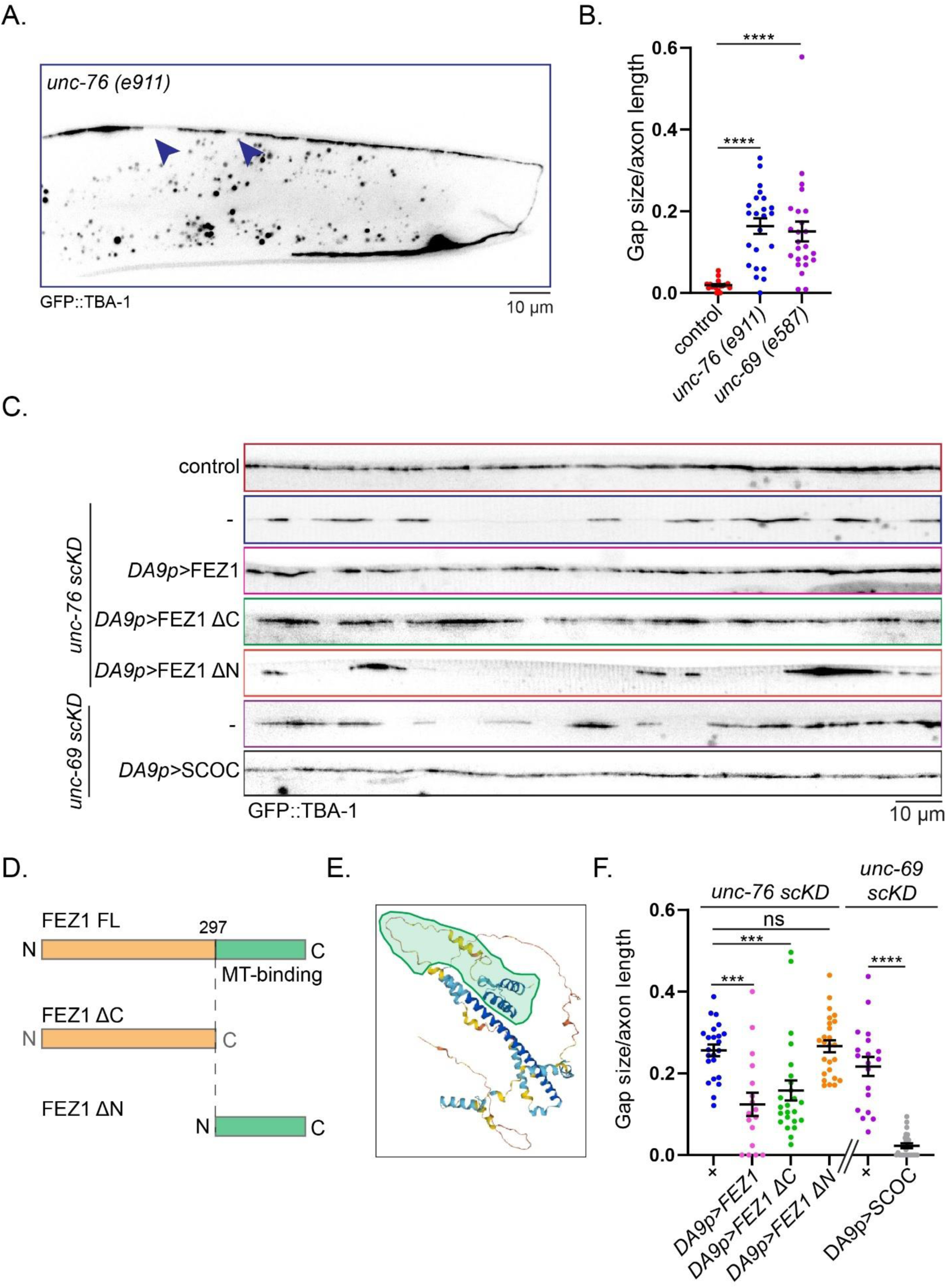
UNC-76/FEZ1 and UNC-69/SCOC function is conserved and FEZ1 MT biding is not required for MT distribution. (A) MT gaps in *unc-76(e911)* hypomorphic mutants. Shown is an example without severe axon guidance defects. Distal is left and proximal right. (B) Quantification of MT gaps in *unc-76(e911)* and *unc-69(e587)* mutants. Control is wildtype. Statistical test is a Kruskall-Wallis test with Dunn’s multiple comparisons, ****=p<0.0001, n=20-24, error bars show SEM. (C) MT gaps in axons of *unc-76 scKD* and *unc-69 scKD* mutant animals with DA9-specific expression of their human homologs FEZ1 and SCOC, respectively. FEZ1 and SCOC are able to rescue MT gaps, and FEZ1 requires its N-ter moiety for this rescue. FEZ1 lacking its C-ter, which contains a putative MT binding domain, can still rescue *unc-76 scKD* mutants. Scale bar=10 µm. (D,E) Cartoon and Alpha-Fold III model of FEZ1, with the C-ter MT binding domain indicated. (F) Quantification of C. MT gaps are normalized to length of the axon. Statistical test for *unc-69 scKD* compared to *unc-69 scKD; DA9p>SCOC* is a Mann-Whitney test, ****=p<0.0001. Statistical test for *unc-76 scKD* genotypes is Kruskall-Wallis test using Dunn’s multiple comparisons test with *unc-76 scKD* as the control column, p=0.0004 for *unc-76 scKD* compared to *unc-76 scKD;* DA9p>FEZ1, p=0.0008 for *unc-76 scKD* compared to *unc-76 scKD;* DA9p>FEZ1 ΔC. ns=not significant. n=17-25, error bars show SEM.

**Figure S2:**
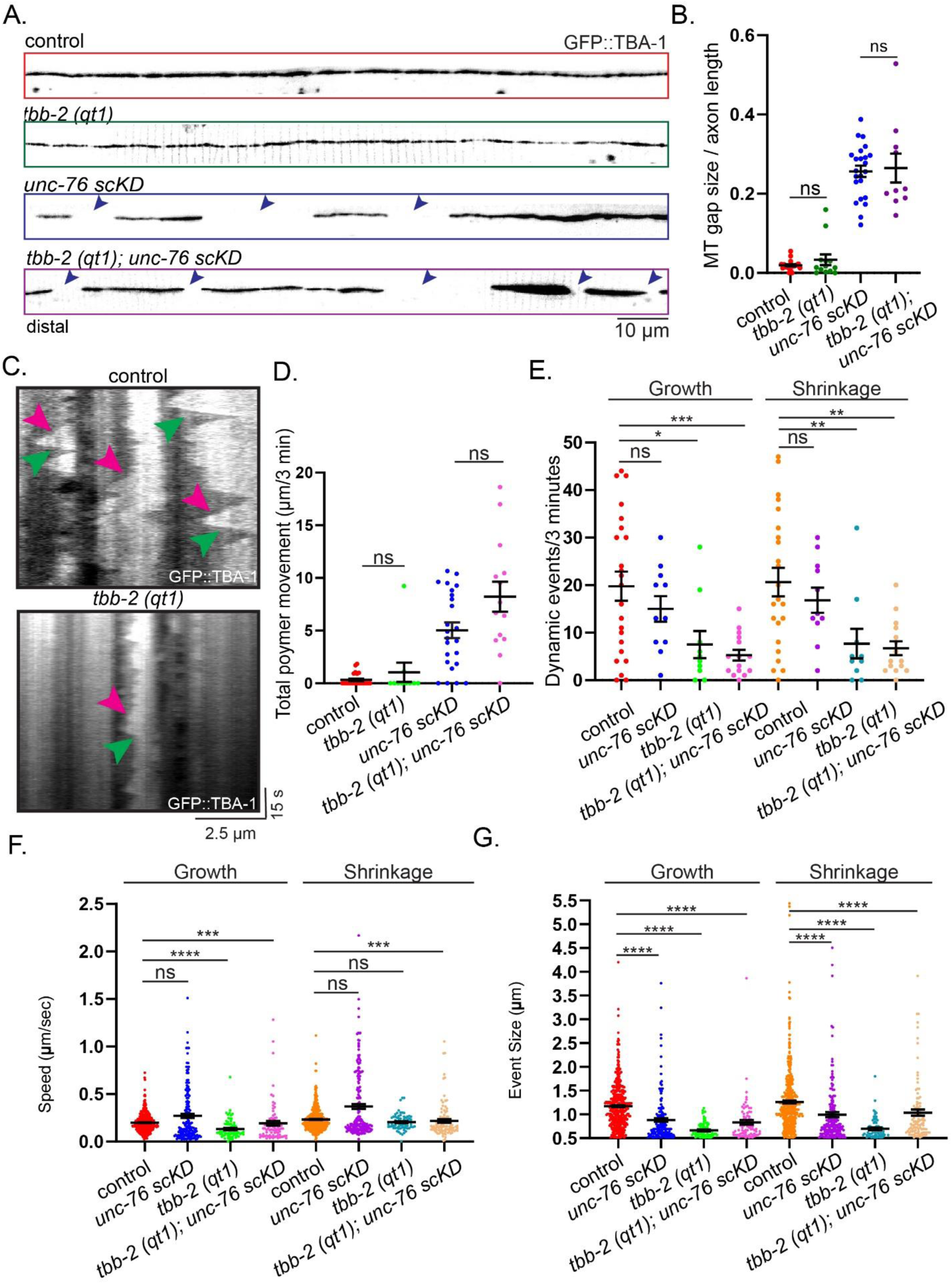
Changes to MT plus-end dynamics are not responsible for the MT gap phenotypes. (A) MT distribution in wildtype and *unc-76 scKD* mutants compared to *tbb-2(qt1)* and *tbb-2(qt1); unc-76 scKD* single and double mutants. Blue arrows highlight MT gaps. Animals are oriented such that distal is on the left and proximal is on the right. Scale bar=10 µm. *tbb-2(qt1)* is a ts mutation that suppresses MT dynamics. Animals were transferred from 15 to 25 degrees as L1 to bypass any potential defects to axon outgrowth. (B) Quantification of A. MT gaps are normalized to length of the axon. Both pairs were compared via the Mann-Whitney test, ns=not significant. n=10-23, error bars show SEM. (C) Representative kymographs highlight the reduced MT dynamics in *tbb-2 (qt1)* mutant animals. MT growth events are highlighted with magenta arrows and MT catastrophe events are highlighted with green arrows. The left side of the kymograph is distal and the right side is proximal. Frame rate=0.333s. (D) Quantification of MT movements in indicated genotypes. *tbb-2(qt1)* does not suppress the MT movements in *unc-76 scKD* mutants. Mann-Whitney tests, ns=not significant, n=10-24, error bars show SEM. (E) Quantification of the number of MT dynamic events with event size>0.5 µm per 3-minute movies. Statistics for growth and shrinkage were calculated via two one-way ANOVAs with Dunnett’s multiple comparisons with wildtype as the control column for each test. *=0.0127, ***=0.0005, ns=not significant for anterograde events. Control compared to *tbb-2 (qt1)*, **=0.0086, control compared to *tbb-2(qt1); unc-76 scKD* **=0.0010 for retrograde events. n=10-23 animals, error bars show SEM. (F) Quantification of the speed of MT dynamic events with event size>0.5 µm were measured over the course of 3 minute movies. Statistics for growth and shrinkage were calculated via two Kruskall-Wallis tests using Dunn’s multiple comparisons test. For anterograde, ****=p<0.0001, ***=p=0.0008, ns=not significant, n=72-455 dynamic events. For retrograde, ***=p=0.0002, ns=not significant, n=73-475 dynamic events measured in n=10-23 animals, error bars show SEM. (G) Quantification of the event size of MT dynamic events with event size>0.5 µm were measured over the course of 3 minute movies. Statistics for growth and shrinkage were calculated via two Kruskall-Wallis tests using Dunn’s multiple comparisons test with wildtype as the control column for each test. For anterograde, ****=p<0.0001, n=72-455 dynamic events in n=10-23 animals. For retrograde, ****=p<0.0001, n=73-475 dynamic events in n=10-23 animals, error

**Figure S3:**
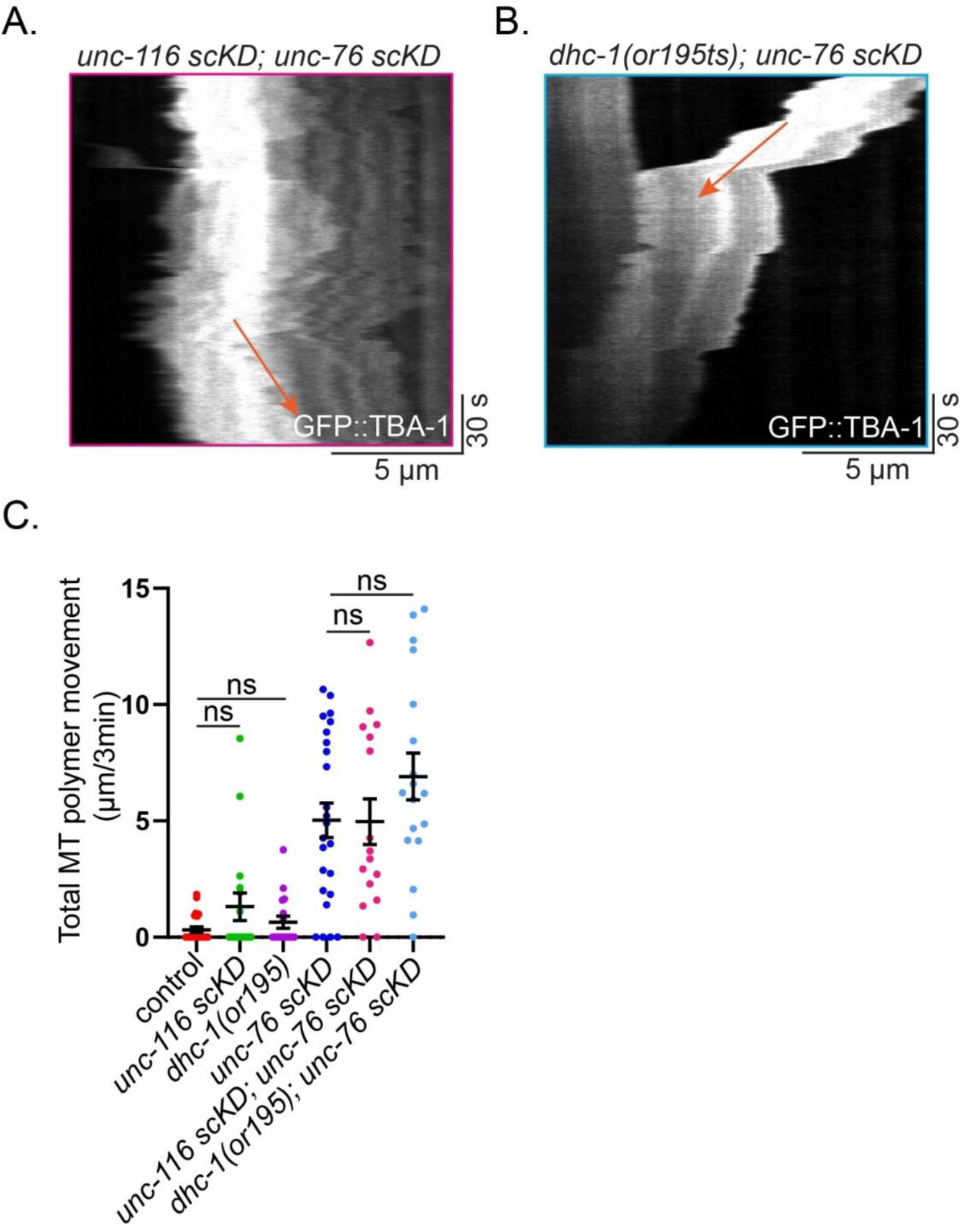
kinesin-1/*unc-116* and dynein heavy chain/*dhc-1* are not required for MT movements in *unc-76 scKD* mutants. (A) Representative kymograph highlights MT movement in *unc-116 scKD; unc-76 scKD* double mutant animals. Orange arrows highlight long, processive MT movements. Frame rate=0.333s. Animals were transferred to auxin containing plates to initiate UNC-116 degradation as L1 larvae, after axon navigation to its target. (B) Representative kymograph highlights MT movement in *dhc-1(or195ts); unc-76 scKD* double mutant animals. Orange arrows highlight long, processive MT movements. Frame rate=0.333s. *dhc-1(or195ts)* is a ts-lethal allele. Animals were grown at 15 degrees and shifted to 25 degrees as L1 larvae to avoid sterility and axon growth defects. (C) Quantification of MT movements over the course of 3-minute movies. Control, *unc-116 scKD*, and *dhc-1(or195ts)* single mutants were compared by Kruskall-Wallis test using Dunn’s multiple comparisons, ns=not significant, n=17-23. *unc-76 scKD; unc-116 scKD; unc-76 scKD,* and *dhc-1 (or195ts); unc-76 scKD* were compared by Kruskall-Wallis test using Dunn’s multiple comparisons, ns=not significant, n=16-24, error bars show SEM.

**Movie S1**: Representative 3-minute movies of wildtype, *unc-76 scKD*, *unc-69 scKD*, and *unc-70 scKD* distal axons. Scale bar represents 10 microns. Movies were acquired at 3 frames per second and are shown at 30 frames per second.

